# Targeting the MYC interaction network in B-cell lymphoma via histone deacetylase 6 inhibition

**DOI:** 10.1101/2021.06.01.445760

**Authors:** René Winkler, Ann-Sophie Mägdefrau, Eva-Maria Piskor, Markus Kleemann, Mandy Beyer, Kevin Linke, Lisa Hansen, Anna-Maria Schaffer, Marina E. Hoffmann, Simon Poepsel, Florian Heyd, Petra Beli, Tarik Möröy, Siavosh Mahboobi, Oliver H. Krämer, Christian Kosan

**Author notes:** These authors contributed equally to this work (René Winkler, Ann-Sophie Mägdefrau). Present address: Josep Carreras Leukaemia Research Institute (IJC), Campus ICO-Germans Trias i Pujol, Badalona, 08916, Spain.

## Abstract

Overexpression of *MYC* is a genuine cancer driver in lymphomas and related to poor prognosis. However, therapeutic targeting of the transcription factor MYC remains challenging. Here, we show that inhibition of the histone deacetylase 6 (HDAC6) using the HDAC6 inhibitor Marbostat-100 (M-100) reduces oncogenic MYC levels and prevents lymphomagenesis in a mouse model of *MYC*-induced aggressive B-cell lymphoma. M-100 specifically alters protein-protein interactions by switching the acetylation state of HDAC6 substrates, such as tubulin. Tubulin facilitates nuclear import of MYC, and MYC-dependent B-cell lymphoma cells rely on continuous import of MYC due to its high turn-over. Acetylation of tubulin impairs this mechanism and enables proteasomal degradation of MYC. M-100 targets almost exclusively B-cell lymphoma cells with high levels of MYC whereas non-tumor cells are not affected. M-100 induces massive apoptosis in human and murine *MYC*-overexpressing B-cell lymphoma cells. We identified the heat-shock protein DNAJA3 as an interactor of tubulin in an acetylation-dependent manner and overexpression of DNAJA3 resulted in a pronounced degradation of MYC. We propose a mechanism by which DNAJA3 associates with hyperacetylated tubulin in the cytoplasm to control MYC turnover. Taken together, our data demonstrate a beneficial role of HDAC6 inhibition in MYC-dependent B-cell lymphoma.

## Introduction

B-cells are prone to lymphomagenesis due to their high proliferative capacity and dependence on physiological DNA damage during V(D)J recombination and affinity maturation in germinal centers (1). Non-Hodgkin’s lymphomas, such as Burkitt’s lymphoma (BL) and diffuse large B-cell lymphoma (DLBCL) are aggressive heterogeneous lymphomas originating from germinal center B-cells (2). BL is typically characterized by translocations of *MYC* to the vicinity of potent immunoglobulin enhancers, such as t(8;14) (1). DLBCL shows various molecular alterations, among them translocations of *BCL6* or *BCL2* (1). However, *MYC* translocations also occur in around 15 % of DLBCL (2), and elevated expression of *MYC* correlates with poor clinical prognosis in B-cell lymphoma (3,4). Overexpression of the transcription factor MYC leads to fatal misregulation of cellular metabolism, cell growth, and signaling pathways (5,6). Moreover, MYC controls proliferation, angiogenesis, and mRNA processing in tumor development (7,8).

MYC is known to associate with microtubules for the nuclear import of MYC (9). Importantly, MYC has a half-life of roughly half an hour when transiently expressed (10). This high turn-over rate and the absence of druggable structures make MYC a difficult target for direct inhibition (11). Attempts to directly target MYC via small molecules did not achieve adequate results as these drugs underwent rapid degradation and showed unfavorably high IC50 values (11). In fact, physiological levels of MYC fulfill crucial functions in many cell types, and pharmacological targeting of MYC should only counteract supraphysiological MYC levels present in malignant cells.

MYC can recruit epigenetic modifiers, such as the histone acetylases p300/CBP or the histone deacetylases (HDACs) HDAC1 and HDAC3 to activate or repress distinct genes in cancer cells (8). Pan-HDAC inhibitors (pan-HDACi) that target several HDACs have been shown to give promising results in hematologic malignancies (12–14). The regulation of non-histone proteins, in particular proto-oncogenes, by HDACi enables the control of many essential cellular processes, such as cell survival, proliferation, protein stability, and protein interactions (15). For example, pan-HDACi treatment has been shown to inhibit BCL-6 function by stabilizing BCL-6 acetylation, which leads to the de-repression of its target genes (16). Interestingly, MYC can be found acetylated at K423 upon pan-HDACi treatment, which decreases *MYC* transcription via autoregulation and results in apoptosis (13). However, the success of pan-HDACi in pre-clinical studies only partially improved treatment for patients with hematological malignancies (17).

The use of HDACi that target singular members of the HDAC family will help to understand separate HDAC functions in hematologic malignancies. For example, treatment of DLBCL with the HDAC6 inhibitor ACY-1215 (Rocilinostat) was shown to activate the unfolded protein response by increasing quantity and acetylation of heat-shock proteins (HSPs), eventually resulting in cell death (18). A first clinical trial involving treatment of lymphoma patients with ACY-1215 was completed and stated a favorable safety profile (19).

Interestingly, HDAC6 represents a microtubule-associated deacetylase and was shown to deacetylate microtubules at K40 (20,21). Recently, a novel HDAC6 inhibitor, Marbostat-100 (M-100), was developed based on the structure of Tubastatin A (22,23). M-100 inhibits the major catalytic domain of HDAC6 with at least 250-fold higher affinity compared to other HDACs and the binding mode was well described (22,23). Moreover, M-100 was well-tolerated in a preclinical model of rheumatoid arthritis and showed favorable pharmacodynamics in mice (22).

Our findings identified the HDAC6 inhibitor M-100 as a new tool to target MYC-dependent lymphomas. M-100 efficiently prevented lymphomagenesis, and induced apoptosis in human and murine lymphoma cells by reducing high MYC levels without harming untransformed B-cells. On a molecular basis, we identified a novel cytoplasmic interaction network formed by tubulin, HDAC6, and HSPs that regulates the protein stability of MYC. Our results suggest that inhibition of HDAC6 can be used to target excessive MYC in cancer.

## Results

### HDAC6 inhibition increases the survival of lymphoma-prone mice

The Eμ-Myc mouse line is a commonly used model for studying the spontaneous formation of B-cell lymphomas due to B-cell-specific overexpression of *MYC*, resembling partially disease phenotypes of BL or DLBCL (24). We isolated lymphoma B-cells from Eμ-Myc mice and treated these cells *ex vivo* with the HDAC6 inhibitor M-100. Interestingly, mouse lymphoma cells showed a dose-dependent reduction of Myc protein levels after M-100 treatment (**Figure 1A**). We also detected a strong acetylation of the HDAC6 substrate tubulin concluding efficient HDAC6 inhibition (**Figure 1A**), which is a striking result as M-100 was originally developed against human HDAC6 (22,23). However, levels of unmodified tubulin remained unaltered (**Supplemental Figure 1**). Besides, Parp-1 cleavage was detected in M-100-treated mouse lymphoma cells, indicating apoptosis induction (**Figure 1A**). To test the effect of M-100 *in vivo*, we applied M-100 at 30 mg/kg as described before (22) via intraperitoneal (i.p.) injection to C57BL/6 mice. M-100 treatment did not affect immune cell populations in mice (**Supplemental Figure 2A–C**), as previously shown (22). Acetylation of tubulin was increased in splenic cells 6 h after injection and declined until 72 h post treatment (**Figure 1B**). These results prompted us to test if continuous treatment with M-100 could prevent lymphoma development in Eμ-Myc mice.

**Figure 1:**
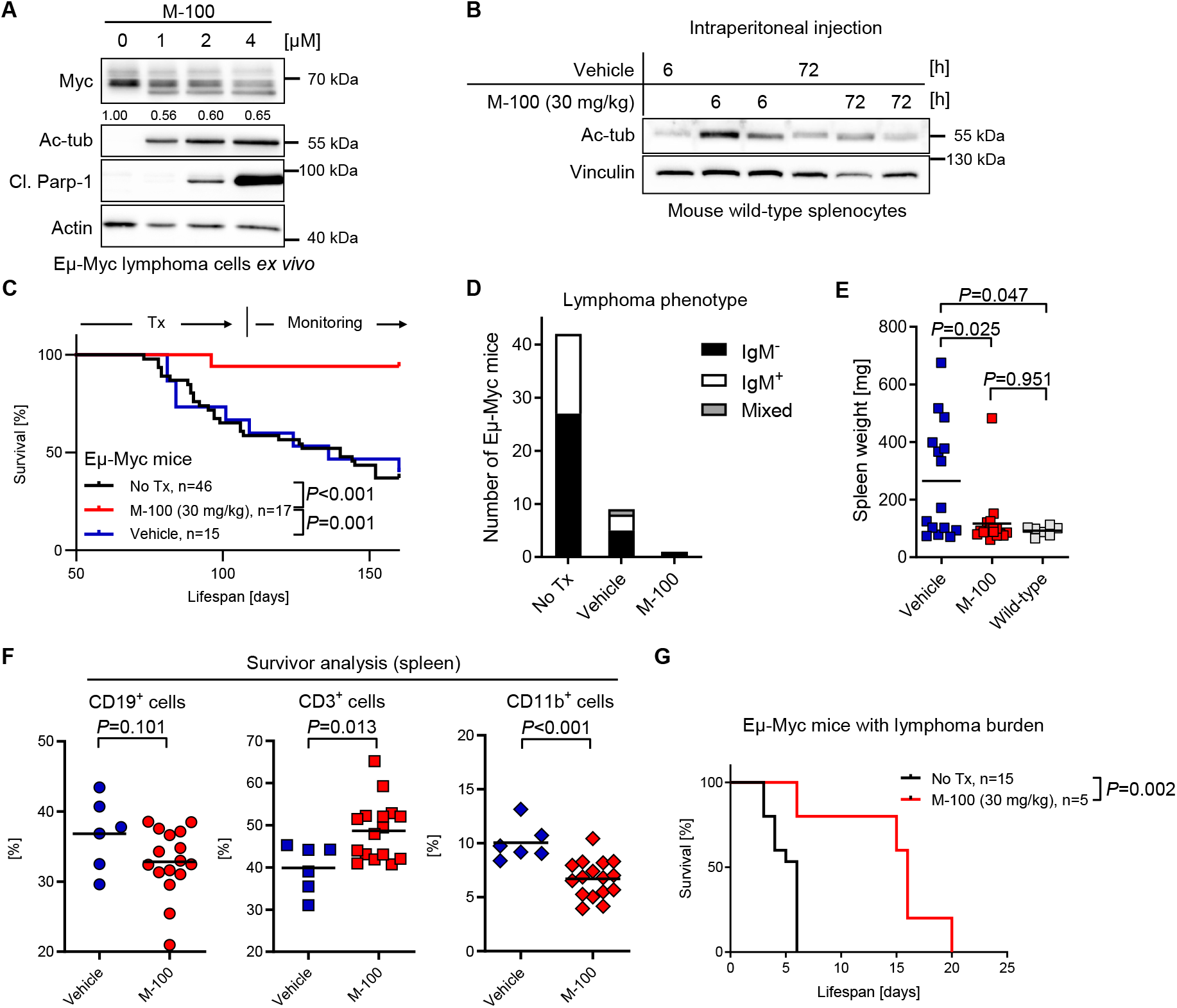
Continuous HDAC6 inhibition increases survival of lymphoma-prone *MYC*-overexpressing mice. **A** Western Blot analysis of Eμ-Myc lymphoma cells isolated from malignant lymph nodes and treated *ex vivo* with increasing concentrations of M-100 for 24 h. Levels of Myc were quantified to untreated conditions. Actin was used as a loading control. Cl. – cleaved. **(B)** Western Blot analysis of splenocytes from wild-type mice treated once via i.p. injection with 30 mg/kg M-100 or vehicle for the indicated time. Each lane represents one individual mouse. Vinculin was used as a loading control. Ac-tub – acetylated tubulin. **(C)** Survival curves of Eμ-Myc mice treated every 72 h with 30 mg/kg M-100 (n=17) or vehicle (n=15). Treatment (Tx) started at age 70 d for six weeks. Mice were monitored for additional six weeks for signs of lymphoma development. Median survival vehicle cohort: 136 d, M-100 cohort: not possible to calculate, untreated Eμ-Myc mice: 140 d. Log-Rank-test. **(D)** Lymphoma phenotypes of diseased Eμ-Myc mice measured by flow cytometry. **(E)** Spleen weights of all mice from the different cohorts. Spleen weights from wild-type mice serve as control. Wilcoxon rank-sum test. **(F)** Flow cytometry analysis of splenic immune cell populations of survivors. Unpaired Student’s t-test, two-tailed. **(G)** Survival curves of Eμ-Myc mice suffering from lymphoma treated with M-100 (median 16 d) or left untreated (median 6 d). Mice received M-100 (30 mg/kg) every 72 h starting when palpable lymph node swelling was present. Treatment continued until endpoint criteria were reached. Log-Rank-test. Data in (A) are representative of n=3 independent experiments. (E) and (F): Each dot represents one mouse. Bars depict mean.

To study the long-term effect of HDAC6 inhibition by M-100 on B-cell lymphomagenesis, 70-day-old Eμ-Myc mice with a high incidence for lymphoma development (25), received i.p. injections with 30 mg/kg M-100 or vehicle every 72 h for six weeks. Mice were monitored for an additional period of six weeks for spontaneous tumor formation after M-100 withdrawal. M-100 treatment of Eμ-Myc mice significantly increased the overall survival compared to vehicle-treated and untreated mice, even after M-100 withdrawal (**Figure 1C**). While 60 % of vehicle-treated Eμ-Myc mice developed lymphomas in this period, only one out of 17 M-100-treated mice manifested a lymphoma (B220^+^ IgM^−^) and had to be sacrificed. Lymphomas from non-treated and vehicle-treated Eμ-Myc mice showed a 2:1 ratio of IgM^−^to IgM^+^ B-cell tumors indicating that lymphoma cells in this model system have an immature or mature B-cell origin (**Figure 1D**), as observed before (26). Long-term treatment with M-100 did not affect the body weight of mice (**Supplemental Figure 2D**). The mean spleen weight of Eμ-Myc mice, however, was significantly decreased after continuous M-100 treatment, indicating reduced disease progression (**Figure 1E, Supplemental Figure 2E, F**). We analyzed immune cell populations of surviving mice by flow cytometry. M-100-treated Eμ-Myc mice showed a slightly reduced splenic CD19^+^ B-cell pool (**Figure 1F, Supplemental Figure 3A**). B-cell development in the bone marrow was despite this not affected by M-100 treatment (**Supplemental Figure 3B**) and increased apoptosis of B-cells was not detected *in vivo* (**Supplemental Figure 3C**). In fact, B220^low^ cells were still present in spleens (**Supplemental Figure 3C**), which have been previously described as pre-tumor B-cells in this model (27,28). Besides, survivors from the M-100 cohort showed a significant decline in CD11b^+^ myeloid cells and an absolute as well as a relative increase in CD3^+^ T-cells compared to the vehicle group (**Figure 1F, Supplemental Figure 3A**). Here, further investigation showed that survival of primary CD4^+^ and CD8^+^ T-cells was not affected by M-100 treatment, whereas proliferation of CD4^+^ T-cells after M-100 treatment was even increased when cultured together with B-cells (**Supplemental Figure 3D–F**). These results suggest that M-100 affects myeloid cell abundance and T-cell proliferation which might have contributed to prevent B-cell lymphomagenesis in Eμ-Myc mice.

Next, we assessed the curative potential of M-100 on already manifested lymphoma. We treated lymphoma-bearing Eμ-Myc mice with M-100 and analyzed their survival. After the detection of lymphoma, untreated Eμ-Myc mice died within 6 days. Acute M-100 treatment of sick Eμ-Myc mice significantly improved survival almost 3-fold, which was also indicated by maintaining lower disease severity scores (**Figure 1G, Supplemental Figure 4**). Taken together, HDAC6 inhibition strongly reduced B-cell lymphomagenesis and extended life span in mice overexpressing *MYC* in the B-cell compartment.

### Inhibition of HDAC6 specifically induces apoptosis in B-cell lymphoma

To test the effects of HDAC6 inhibition on lymphoma B-cells *ex vivo*, purified lymphoma cells from Eμ-Myc mice were treated for 72 h with increasing amounts of M-100. Purified B-cells from wild-type mice were activated with 10 μg/ml LPS and served as control. B-cells from Eμ-Myc mice showed a strong induction of apoptosis already after treatment with 1 μM M-100 (**Figure 2A**). Of note, activated B-cells from wild-type mice did not show an increased apoptosis rate (**Figure 2A**). We also assessed the effect of M-100 on the cell cycle of lymphoma and activated wild-type B-cells. Murine lymphoma cell populations showed a significant increase of cells in the subG1 fraction, indicating apoptosis, and a reduction in G1/S-phases 72 h after M-100 treatment (**Figure 2B, C**). However, non-malignant B-cells did not any show cell cycle alterations (**Figure 2B, C**). Gene expression of *Bbc3* and *Pmaip1*, encoding pro-apoptotic Puma and Noxa was increased in lymphoma but not wild-type B-cells after M-100 treatment (**Figure 2D**). Conversely, transcription of *Bcl2* was strongly decreased in lymphoma cells after HDAC6 inhibition (**Figure 2D**), which underlines the apoptotic phenotype (**Figure 2A–C**). M-100 also affected Myc protein levels in activated wild-type B-cells that was not caused by transcriptional changes (**Figure 2D, E**). However, wild-type B-cells showed a strong upregulation of the anti-apoptotic factor Bcl-2 after M-100 treatment that was not present in Eμ-Myc cells, and which likely protected wild-type B-cells from apoptosis induction (**Figure 2E**). To further examine the role of Bcl-2, we treated proliferating B-cells from wild-type mice with the highly specific BCL-2 inhibitor Venetoclax (**Figure 2F**). Inhibition of Bcl-2 in combination with M-100 led to a strong apoptosis induction of activated B-cells, which indicates that upregulation of Bcl-2 mediated survival signals in healthy B-cells (**Figure 2F**). We also checked the effects of M-100 on CH12F3 cells, a murine B-cell line harboring no *MYC* translocation (29). In line with the findings from wild-type B-cells, no apoptosis or altered cell cycle was measured after M-100 treatment in CH12F3 cells (**Figure 2G**). Taken together, M-100 exclusively induced apoptosis in murine lymphoma cells with elevated Myc levels as these cells failed to upregulate Bcl-2.

**Figure 2:**
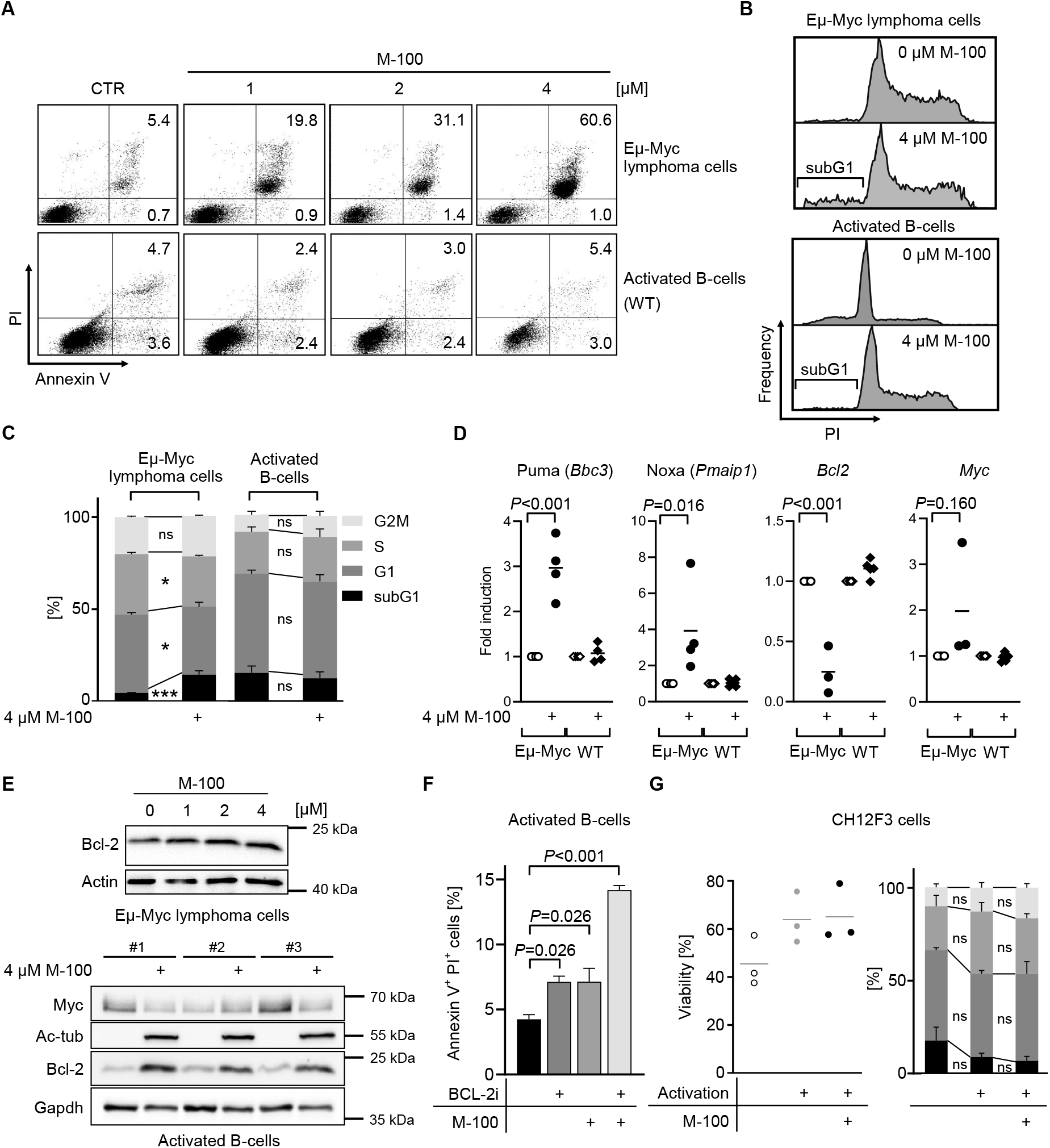
M-100 specifically induces apoptosis in murine lymphoma B-cells but not activated wild-type B-cells. **A** Apoptosis detection using Annexin V and PI staining. Eμ-Myc lymphoma and activated (10 μg LPS/ml) wild-type mouse B-cells were compared. **(B)** Representative cell cycle analysis of Eμ-Myc lymphoma and activated B-cells treated for 72 h. **(C)** Cell cycle analysis from (B) was quantified. Two-Way ANOVA (Sidak’s posthoc). **(D)** Gene expression changes of Eμ-Myc lymphoma or activated wild-type (WT) B-cells cells treated for 24 h with 4 μM M-100 using quantitative real-time PCR analysis. One-Way ANOVA (Tukey’s posthoc). **(E)** Western Blot analysis of activated wild-type B-cells or Eμ-Myc lymphoma cells treated with M-100 for 24 h. Gapdh and actin were used as loading controls. Ac-tub – acetylated tubulin. **(F)** Apoptosis was analyzed in activated wild-type B-cells treated with 4 μM M-100, 0.5 μM BCL-2 inhibitor (BCL-2i) Venetoclax or both for 24 h. One-Way ANOVA (Tukey’s posthoc). **(G)** CH12F3 cells were analyzed regarding viability (Annexin V/PI staining) and cell cycle after treatment with 2 μM M-100 for 48 h. Activation was performed using 1 μg/ml CD40L, 5 ng/ml IL-4, and 1 ng/ml TGF-β and added 2 h after M-100 treatment. CH12F3 cells harbor no *MYC* translocation. Two-Way ANOVA (Sidak’s posthoc). Data in (A) – (G) are representative of at least n=3 independent experiments. Data represent mean + SEM, if applicable. \**P*<0.05, \*\*\**P*<0.001, ns – not significant.

### HDAC6 inhibition results in apoptosis of human B-cell lymphoma cells

We also assessed the response to M-100 in human B-cell lymphoma cell lines of the BL and DLBCL type characterized by *MYC* translocation or amplification (30–35), although the individual *MYC* mutation profile differed greatly in these cells (36,37), (**Table 1**). We determined the efficacy of M-100 by MTT assay, and four out of five tested cell lines showed striking dose-responses to M-100 (**Figure 3A**). The obtained IC50 values ranged between 2.07 μM and 5.25 μM. Raji cells, however, were less sensitive to M-100 (IC50=17.17 μM), although a moderate apoptosis induction was achieved at 4 μM (**Figure 3B**). M-100 treatment resulted in the induction of apoptosis in all tested human BL and DLBCL cell lines as demonstrated by the accumulation of Annexin V^+^ cells (**Figure 3B**). Consistent with our findings from murine lymphoma cells, we detected a significant increase of subG1 fractions 48 h post M-100 treatment (**Figure 3C**), and a reduced entry into S-phase already 24 h post treatment in human lymphoma cells (**Supplemental Figure 5A–C**). Only BL-30 cells showed an unaltered cell cycle profile, which might be explained by the high mutation rate within the *MYC* gene (**Figure 3B,Table 1**). To confirm apoptosis as the responsible cell death mechanism in Ramos cells, combinatorial treatment of M-100 and the caspase-specific inhibitor Z-VAD-FMK was performed (**Supplemental Figure 5D**). The use of both Z-VAD-FMK and M-100 prevented PARP-1 and Caspase 3 cleavage and ultimately reduced the number of early apoptotic cells (**Supplemental Figure 5E**). These results clearly show that cell death induction by M-100 is facilitated via the apoptotic pathway.

**Table 1:** *MYC* mutations in human B-cell lymphoma cell lines. Table summarizing *MYC* mutations in the used cell lines from the CCLE and COSMIC databases. °Sequencing data were only available for subclone Ramos2G64C10. BL – Burkitt’s lymphoma, DLBCL – diffuse large B-cell lymphoma.

| Cell line | MYC mutations | Type |
| --- | --- | --- |
| Ramos° | Translocated | BL |
| Raji | Translocated<br>p.YQ32del, p.S6T, p.E39D, p.A44V,<br>**p.T58I** | BL |
| BL-30 | Translocated<br>p.Q50H, p.T73I, p.G105D, p.S218N | BL |
| SUDHL-6 | Translocated | DLBCL |
| OCI-Ly3 | Amplified | DLBCL |

**Figure 3:**
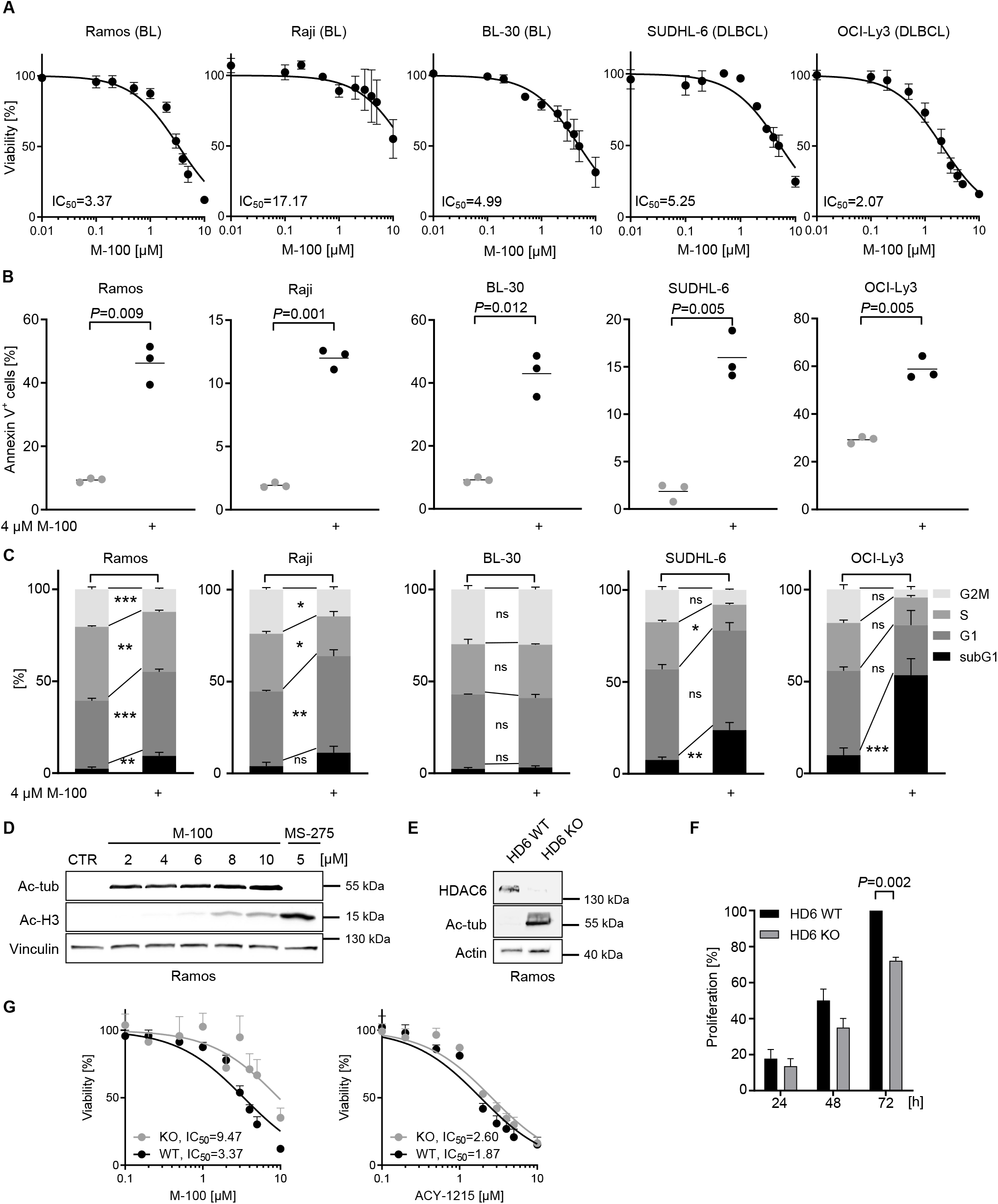
Different human B-cell lymphoma cell lines respond to HDAC6 inhibition by initiating apoptosis and cell cycle arrest. **A** Cell viability of human B-cell lymphoma cells after treatment with increasing concentrations of M-100 for 48 h using MTT assay. Non-linear regression (inhibitor vs. normalized response) was inserted. **(B)** Amount of Annexin V^+^ cells after treatment with 4 μM M-100 for 48 h. Unpaired Welch’s t-test, two-tailed. **(C)** Cell cycle analysis of cells treated with 4 μM M-100 for 48 h. Two-Way ANOVA (Sidak’s posthoc). **(D)** Different concentrations of M-100 were tested for inducing ac-H3 signals by Western blot. Vinculin was used as a loading control. **(E)** Ramos HDAC6 (HD6) knock-out (KO) cells were generated and compared to HDAC6 wild-type (WT) cells. Western Blot analysis shows absence of HDAC6. Actin serves as a loading control. **(F)** Proliferation was measured of Ramos HDAC6 WT and HDAC6 KO cells by cell counting, and normalized to WT cells at t=72 h. Two-Way ANOVA (Sidak’s posthoc). **(G)** Dose-response curves were determined for Ramos HDAC6 WT and KO cells treated for 48 h with increasing concentrations of M-100 or ACY-1215 by MTT assay. Non-linear regression (inhibitor vs. normalized response) was inserted. Data in (A) – (G) are representative of at least n=3 independent experiments. Data represent mean + SEM, if applicable. \**P*<0.05, ** *P*<0.01, \*\*\**P*<0.001, ns – not significant.

To exclude off-target effects of M-100, we compared different concentrations of M-100 to the pan-HDACi MS-275 that inhibits HDAC1, HDAC2, and HDAC3. H3 acetylation was induced at concentrations greater than 6 μM M-100 (**Figure 3D**). Importantly, this limit is above most obtained IC50 values for M-100 in B-cell lymphoma cells, suggesting a specific effect of M-100 (**Figure 3A**). Furthermore, M-100 had superior targeting properties compared to the HDAC6 inhibitor ACY-1215 as HDAC6 was efficiently inhibited by M-100 but HDACs that deacetylate histone H3 were not (**Supplemental Figure 6A**). To further validate our findings, we generated Ramos HDAC6 knock-out (KO) cells using targeted CRISPR/Cas9 technology. HDAC6 KO cells were characterized by permanent hyperacetylation of tubulin (**Figure 3E**). Importantly, KO of HDAC6 mimics to some extent the treatment with M-100 as Ramos HDAC6 KO cells showed reduced proliferation and increased apoptosis compared to WT cells (**Figure 3F, Supplemental Figure 6B**). As expected, Ramos HDAC6 KO cells were rather insensitive to M-100 treatment (IC50=9.47 μM) as measured with MTT assay and apoptosis detection by Annexin V/PI staining (**Figure 3G, Supplemental Figure 6B**). However, KO of HDAC6 barely altered the response of Ramos cells towards ACY-1215, revealing a very narrow window of specific targeting (**Figure 3G, Supplemental Figure 6B**). This underlines that M-100 has improved specificity for HDAC6 compared to ACY-1215 whose off-targets were recently described in detail (38).

### HDAC6 inhibition causes MYC degradation

Human *MYC*-overexpressing lymphoma cells responded to M-100 treatment with a strong induction of apoptosis. To shed further light on the underlying cellular mechanism, we treated Ramos cells with different concentrations of M-100 for 6 h and 24 h. MYC protein levels were already reduced 6 h after treatment with 2 μM and 4 μM M-100 (**Figure 4A, Supplemental Figure 7A**). Consistent with our previous findings from murine lymphoma cells, we also detected PARP-1 cleavage 24 h after treatment with M-100 (**Figure 4A**). On the contrary, BCL-2 levels decreased after HDAC6 inhibition although transcription of the *BCL2* gene was elevated, assuming a possible feedback loop (**Figure 4A, Supplemental Figure 7B, C**). Expression of *BIM*, encoding a pro-apoptotic mediator, was significantly upregulated which was also present at the protein level (**Supplementary Figure 7B, C**). As in murine lymphoma cells, the decrease of MYC was not due to transcriptional changes (**Supplementary Figure 7C**). To determine if MYC degradation is regulated by ubiquitin-mediated proteolysis upon HDAC6 inhibition, we performed combinatorial treatments with M-100 and the proteasome inhibitor MG132 (**Figure 4B**). MYC degradation was efficiently blocked when proteasomes were not functioning. The use of MG132 resulted in pronounced general accumulation of ubiquitinated proteins, in contrast to M-100 (**Figure 4C**). In addition, immunoprecipitation (IP) of MYC after HDAC6 inhibition revealed ubiquitinated MYC protein bands at molecular weights of 85 kDa and 100 kDa (**Figure 4C**). In fact, all tested BL and DLBCL cell lines showed a substantial decrease of MYC protein after 6 h treatment with M-100 (**Figure 4D**), pointing towards a general mechanism of MYC degradation after HDAC6 inhibition. We also observed a slight increase in MYC turnover when HDAC6 was knocked out (**Supplementary Figure 8**). These results indicate that HDAC6 inhibition by M-100 specifically provokes ubiquitination of MYC and subsequent proteasomal degradation.

**Figure 4:**
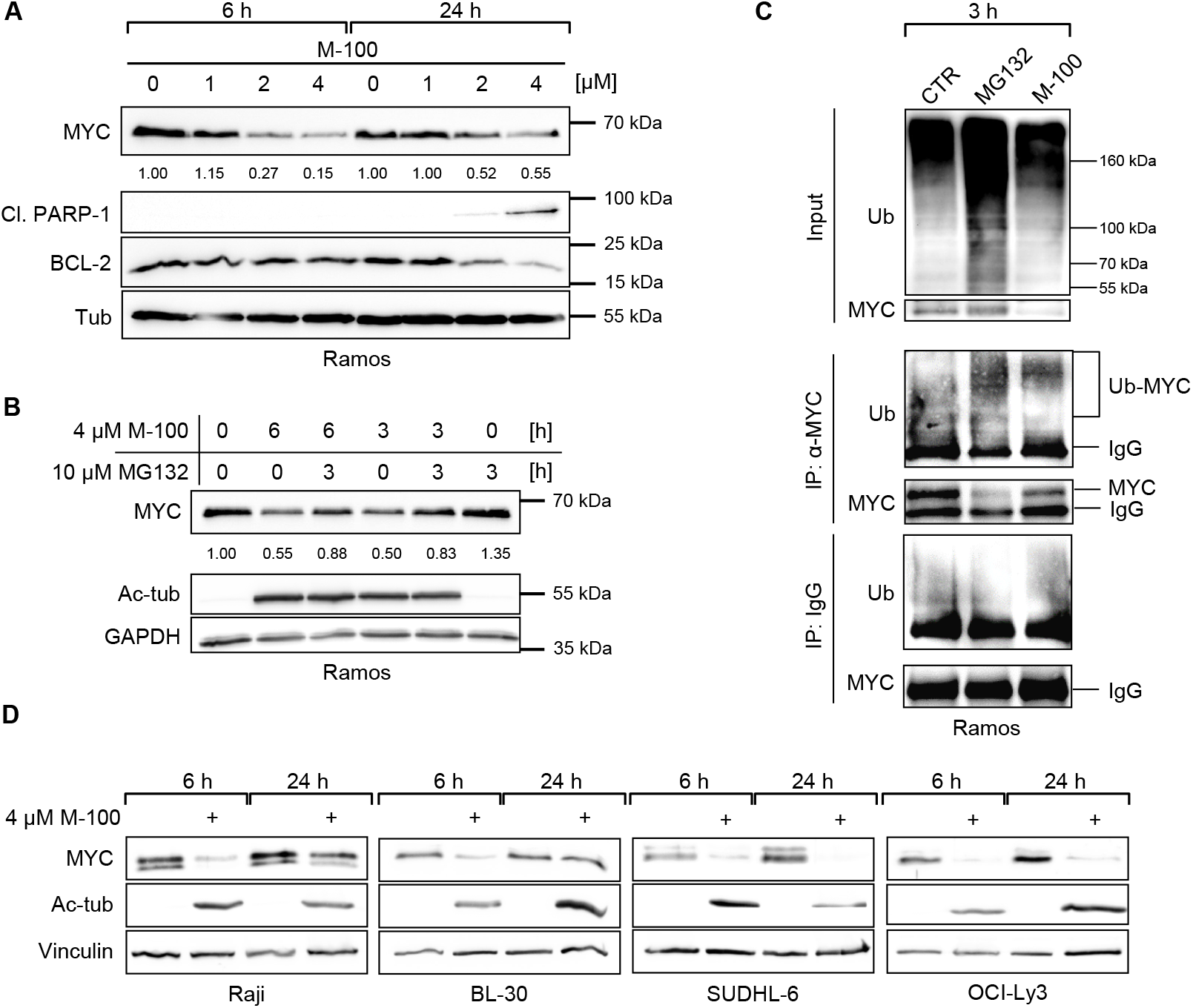
HDAC6 inhibition results in rapid *MYC* degradation. **A** Western Blot analysis of Ramos cells treated for the indicated time with different concentrations of M-100. Levels of MYC were quantified to untreated conditions. Tubulin was used as a loading control. Cl. – cleaved. **(B)** Western Blot analysis of Ramos cells treated with 10 μM MG132 to block proteasomal degradation and/or 4 μM M-100 for the indicated time. Levels of MYC were quantified to untreated conditions. GAPDH was used as a loading control. **(C)** Ubiquitination of proteins was analyzed in Ramos cells treated for 3 h with 10 μM MG132, 4 μM M-100 or left untreated by Western blot. 50 μg protein was loaded for input. Endogenous MYC was immunoprecipitated from these cell lysates. Unspecific IgG was used for control IPs. Ub – Ubiquitin. **(D)** Western Blot analysis of B-cell lymphoma cell lines treated for 6 h and 24 h with 4 μM M-100. Vinculin was used as a loading control. Ac-tub – acetylated tubulin. Data in (A) – (D) are representative of n=3 independent experiments.

### MYC degradation is associated with changes in the interactome of hyperacetylated tubulin

To test whether the proteasomal degradation of MYC is mediated by a direct interaction, we performed IPs and were able to pull-down endogenous HDAC6/MYC complexes in Ramos cells (**Figure 5A**). These complexes also appeared when the catalytic activity of HDAC6 was inhibited. However, HDAC6 is localized exclusively in the cytoplasm, whereas MYC is found predominantly in the nucleus (**Figure 5B**). Thus, we analyzed the impact of MYC localization for its degradation. For this purpose, we transfected NIH-3T3 cells to express GFP-MYC and treated these cells with Importazole or Leptomycin B to block nuclear import or export, respectively (**Figure 5C**). Afterward, cells were treated with cycloheximide (CHX) to induce cell-intrinsic degradation and levels of MYC-GFP were tracked via flow cytometry. Interestingly, the degradation of MYC-GFP was significantly accelerated in the cytoplasm but not in the nucleus (**Figure 5C**).

**Figure 5:**
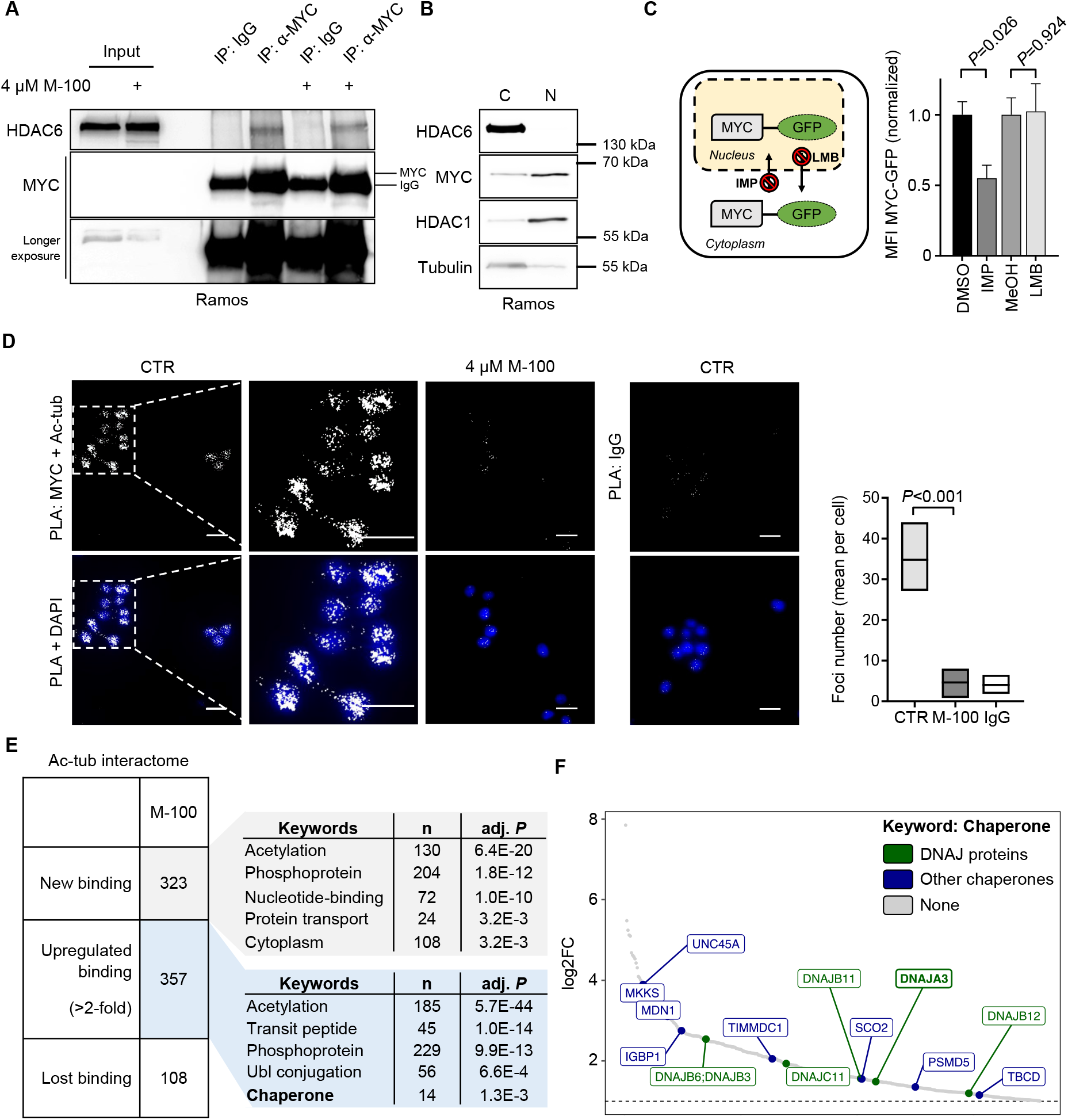
Cytoplasmic MYC degradation is associated with changes in the interactome of hyperacetylated tubulin. **A** Interaction of MYC and HDAC6 detected by IP. Ramos cells were treated for 24 h with 4 μM M-100 or left untreated. IPs with unspecific IgG were used as control. **(B)** <u>C</u>ytoplasmic and <u>n</u>uclear fractions were prepared from Ramos cells. HDAC1 was used as a nuclear marker and tubulin as a cytoplasmic marker. **(C)** NIH-3T3 cells overexpressing MYC-GFP were challenged for 1 h with inhibitors of nuclear import (Importazole, IMP; 40 μM), nuclear export (Leptomycin B, LMB; 20 ng/ml) or solvent. Next, cells were treated for 90 min with CHX (50 μg/ml) before flow cytometry. Living cells were gated using FSC/SSC and normalized median fluorescence intensity (MFI) of MYC-GFP was calculated. Data represent mean + SEM. Unpaired Student’s t-test, two-tailed. **(D)** PLA was performed to detect endogenous co-localization of MYC and acetylated tubulin (actub) in Ramos cells. Cells were treated with 4 μM M-100 for 24 h. Staining with unspecific IgG was used as a control. DAPI was used to stain nuclei. Scale bars indicate 20 μm, magnification 60x. PLA foci were counted and compared. Boxplots depict medium and min to max. One-Way ANOVA (Tukey’s posthoc). **(E)** Global interactome analysis was carried out of immunoprecipitated ac-tub via mass spectrometry. MV4-11 cells were treated for 24 h with 0.5 μM M-100. Shown are counts of proteins with new or increased (>2-fold) binding to ac-tub, or loss of binding after treatment compared to control and IgG binding. Uniprot (UP) keyword annotation was performed with DAVID using all proteins that bound to ac-tub after M-100 treatment. Adjusted (adj.) *P*-values are given. Ubl – Ubiquitin-like. **(F)** All proteins belonging to the keyword “chaperone” are depicted with their corresponding log2 fold change (FC). DNAJ proteins are marked in green. Data in (A) and (C) are representative of n=3 independent experiments. Data in (B) and (D) are representative of n=2 independent experiments.

We wanted to analyze the localization of MYC in response to HDAC6 inhibition by M-100. For this purpose, we treated Ramos cells that harbor very little cytoplasmic compartment with M-100 and used proximity ligation assay (PLA). We detected a close localization between MYC and acetylated tubulin under physiological conditions where a small amount of acetylated tubulin is present (**Figure 5D**). However, PLA signals disappeared after M-100 treatment when tubulin was hyperacetylated (**Figure 5D**). It had been previously described that MYC associates with tubulin (9) and our results indicate that this MYC-tubulin interaction is acetylation-dependent.

To further investigate the effect of hyperacetylated tubulin, a detailed interactome analysis of hyperacetylated tubulin was performed in MYC-dependent blood cancer cells via mass spectrometry. Bound proteins were identified and considered by either increased abundance (>2-fold) or new binding to tubulin after M-100 treatment compared to control (**Supplementary Table 1**). From 2309 identified proteins, treatment with M-100 led to enhanced binding of 357 proteins and new binding of 323 proteins to hyperacetylated tubulin (**Figure 5E**). A functional annotation of the altered tubulin interactome after M-100 treatment using the DAVID tool confirmed significant changes in proteins related to acetylation, nucleotide-binding, protein transport, and ubiquitin conjugation (**Figure 5E**). Overrepresented protein groups that attached to acetylated tubulin after M-100 treatment were HSPs from the chaperone type, including DNAJ proteins (**Figure 5F**). For example, the chaperone member DNAJA3 occurred which is important for proteasomal degradation and was shown to interact with MYC in high-throughput studies (39). Taken together, we demonstrate that MYC degradation is associated with changes in the interactome of heavily acetylated tubulin.

### The chaperone DNAJA3 is recruited to acetylated tubulin and induces MYC degradation

We verified the highly increased binding of DNAJA3 to acetylated tubulin after M-100 treatment using IP techniques and overexpression of DNAJA3-Flag (**Figure 6A, Supplemental Figure 9A**). In addition, we demonstrated the endogenous presence of acetylated tubulin and DNAJA3 complexes in Ramos cells (**Figure 6B, Supplemental Figure 9B**). Of note, the chaperone DNAJA3 has large and small isoforms that are generated from the cleavage of precursor proteins (40,41), which we could also detect in our experiments (**Figure 6A, B**). We used PLA to uncover a close cellular localization of DNAJA3 with MYC in B-cell lymphoma cells (**Figure 6C**). However, HDAC6 inhibition rapidly reduced the number of detected PLA foci per cell. These data suggest that tubulin acts as an interaction hub for MYC, HDAC6 and DNAJA3 in the cytoplasm, and switching the acetylation state of tubulin to hyperacetylation results in disappearance of MYC.

**Figure 6:**
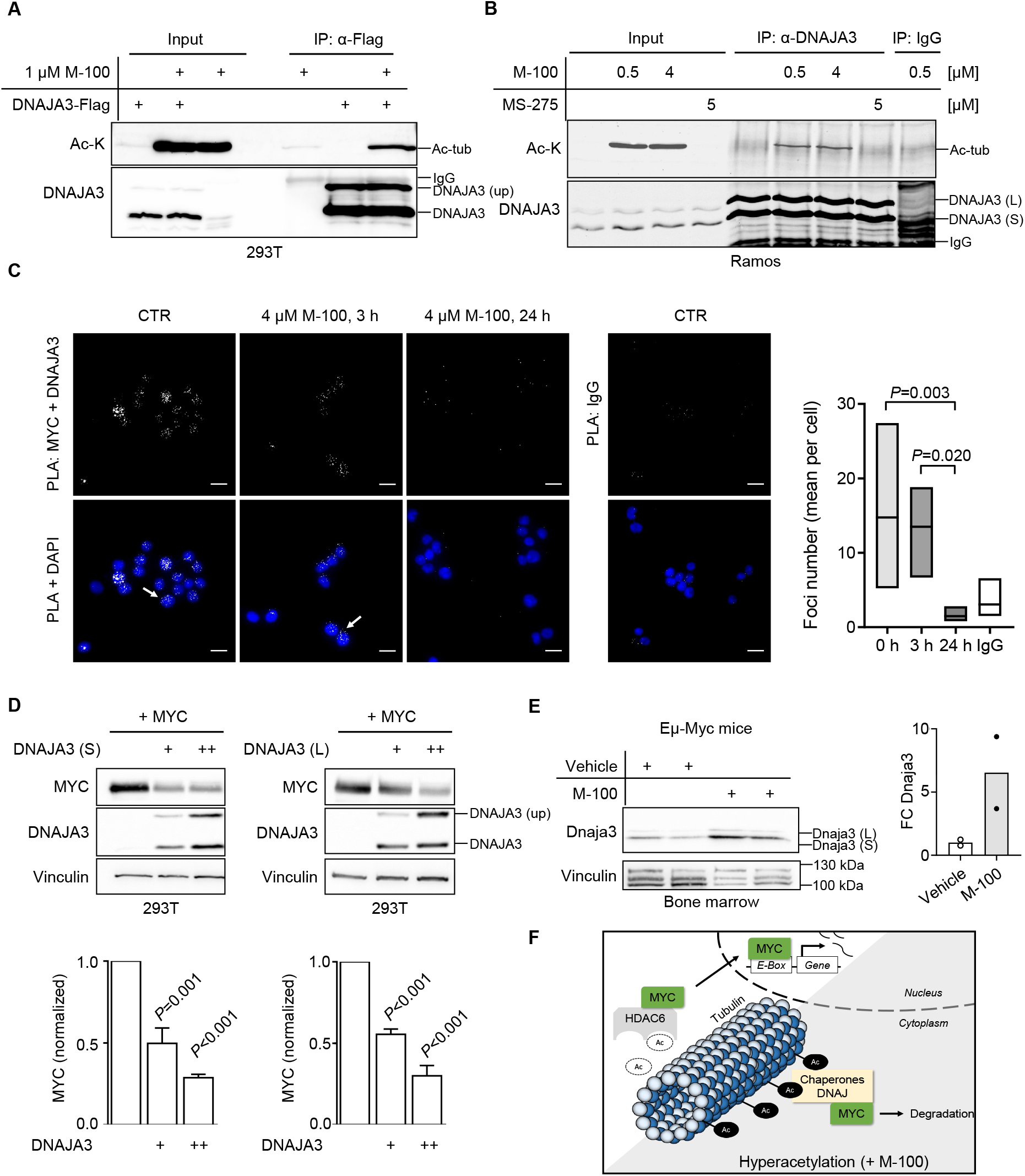
The heat-shock protein DNAJA3 is recruited to hyperacetylated tubulin and induces MYC degradation. **A** Interaction of acetylated tubulin (ac-tub) and DNAJA3. 293T cells were transfected with plasmids encoding DNAJA3-Flag and treated with 1 μM M-100 for 24 h. Cells were lysed in stringent lysis buffer containing M-100. Lysates were used for IP with α-Flag antibodies to precipitate DNAJA3-Flag. Overexpression of DNAJA3 generates unprocessed (up) precursor proteins. **(B)** Endogenous interaction of acetylated tubulin and DNAJA3 in Ramos cells. Cells were treated for 24 h with either 0.5 μM, 4 μM M-100, 5 μM MS-275, or left untreated. Lysates were used for IP with α-DNAJA3 antibodies and tested for interaction with ac-tub. IPs with unspecific IgG were used as control. **(C)** PLA was performed to detect endogenous co-localization of MYC and DNAJA3 in Ramos cells. Cells were treated with 4 μM M-100 for the indicated time points. Staining with unspecific IgG was used as a control. DAPI was used to stain nuclei. Scale bars indicate 20 μm, magnification 60x. PLA foci were counted and compared. Boxplots depict medium and min to max. One-Way ANOVA (Tukey’s posthoc). **(D)** Western Blot analysis of 293T cells overexpressing MYC and increasing amounts of small (S) or large (L) isoforms of DNAJA3. Vinculin was used as a loading control. Quantification of MYC was performed based on Vinculin. One-Way ANOVA (Dunnett’s posthoc). **(E)** Western Blot analysis of bone marrow lysates from Eμ-Myc mice after one i.p. injection with M-100 (30 mg/kg) or vehicle. Small and large isoforms of Dnaja3 can be noticed. Vinculin was used as a loading control. Each lane represents one individual mouse. Quantification of Dnaja3 protein levels is shown based on Vinculin. FC – fold change. **(F)** Scheme summarizing our findings. Hyperacetylation of tubulin by HDAC6 inhibition results in the recruitment of chaperone complexes including the HSP DNAJA3. High levels of DNAJA3 induce degradation of MYC preventing cancer-specific gene regulation. Data in (A), (B), and (D) are representative of n=3 independent experiments, data in (C) are representative of n=2 independent experiments. Data represent mean + SEM, if applicable.

As DNAJ proteins are involved in ATP-dependent protein folding and degradation (42), we investigated potential effects of DNAJA3 on MYC stability. Surprisingly, both DNAJA3 isoforms were able to significantly decrease high MYC levels in overexpression studies in a dose-dependent manner (**Figure 6D**). We retrospectively investigated the bone marrow of Eμ-Myc mice where *MYC*-overexpressing B-cells develop. When Eμ-Myc or wild-type mice were treated once with 30 mg/kg M-100, we were able to detect elevated levels of Dnaja3 (**Figure 6E, Supplementary Figure 10**). Increased levels of Dnaja3 after M-100 treatment might further explain the absence of lymphomagenesis in Eμ-Myc mice.

Taken together, our findings indicate that HDAC6 inhibition results in (1) a remodeling of the tubulin interactome driven by hyperacetylation, (2) recruitment of HSPs including DNAJA3 to acetylated tubulin, and (3) degradation of excessive MYC in *MYC*-overexpressing cells leading to apoptosis (**Figure 6F**). Importantly, our results could be of beneficial use for the therapy of human MYC-dependent lymphoid malignancies.

## Discussion

The role of HDAC6 inhibitors in cancer therapy is still a matter of debate. Recent studies showed that many cancer models are tolerant to HDAC6 inhibitor treatment (43–45). However, a deeper look into the different tumor models considering newly emerging inhibitors might be helpful in developing new strategies for cancer treatment. Here, we demonstrated that MYC-dependent lymphomas are extremely sensitive to the highly specific HDAC6 inhibitor M-100.

In our work, we reveal that M-100 treatment of B-cell lymphoma cells induces proteasomal degradation of MYC. Besides, we prove that MYC forms a complex with the HSP DNAJA3 and hyperacetylation of tubulin strongly recruits DNAJA3. Studies showed that DNAJA3 associates directly with the E3 ligases HUWE1 and von Hippel-Lindau tumor suppressor (46). HUWE1 is a major E3 ligase for MYC (47), which would connect DNAJA3 to the turnover of MYC. Besides, DNAJA3 was shown to mediate ubiquitination and degradation of oncogenic epidermal growth factor receptor (40). Previous strategies to inhibit HDAC6 using ACY-1215 in B-cell lymphoma resulted in an activation of the unfolded protein response by increasing quantity and acetylation of HSPs (18). Similarly, we observed increased levels of the HSP DNAJA3 in the bone marrow of mice treated with M-100. In addition, our interactome analysis suggests that HDAC6 inhibition leads to the recruitment of several chaperones to acetylated tubulin, which might involve other HSPs in the observed MYC instability as well.

The turn-over of MYC is depending on two opposing phosphorylation events of MYC at T58 and S62 which determines protein stability as a phosphodegron (48,49). Our data show that Raji cells, which are mutated at T58, were the least sensitive to M-100. Thus, efficient MYC degradation after HDAC6 inhibition might require wild-type T58 in MYC. Interestingly, phosphorylation of MYC impeded its interaction with tubulin, resulting in increased MYC stability in BL (48). Conversely, our data indicate that the association of MYC with acetylated tubulin decreases MYC stability. Moreover, our interactome data suggest that hyperacetylation of tubulin leads to recruitment of proteins related to the functional annotation terms “Phosphoprotein”, “Nucleotide-binding”, and “Ubiquitin-like conjugation” which may influence the phosphodegron of MYC. It is likely that microtubule acetylation prolongs cytoplasmic retention of MYC making it accessible for proteasomal degradation after interaction with HSPs, such as DNAJA3.

We did not discover direct effects of M-100 on the transcription of the gene encoding MYC neither in murine nor in human cells. This observation is in contrast to the effects of the pan-HDACi MS-275 in hematological malignancies, which impaired transcription of the *MYC* gene itself (13). It is underlined by several studies that M-100 treatment does not result in altered histone acetylation (22,50,51).

The occupation of MYC on distinct sets of target genes was described to depend on the amount of MYC molecules (5), which might explain the adverse Bcl-2 response of murine healthy and *MYC*-transformed B-cells to M-100 in our experiments. Eμ-Myc lymphoma cells treated with M-100 de-repressed *Bbc3* as well as *Pmaip1* and repressed *Bcl2* expression. Importantly, direct transcriptional regulation of *Bbc3* and *Pmaip1* by MYC was shown to be possible (52,53), and deficiency of *Bbc3* and *Pmaip1* accelerated lymphomagenesis in Eμ-Myc mice (54). Thus, permanent upregulation of *Bbc3* and *Pmaip1* caused by HDAC6 inhibition might be another explanation for the extended survival of Eμ-Myc mice upon M-100 treatment.

Taken together, M-100 has potent anti-tumoral activity by targeting the stability of MYC. A fully water-soluble derivative of M-100 exists, which extends possible *in vivo* use (23). However, drug resistance to HDAC6 inhibitors was described (55), and future directions should aim for rational drug combinations to treat distinct malignancies.

## Materials and methods

*Please refer to Supplemental Methods for standard biochemical or molecular techniques*.

### *In vivo* animal studies

Female and male C57BL/6JRj and Tg(IghMyc)22Bri (“Eμ-Myc”) mice with C57BL/6JRj background were housed in individually ventilated cages in groups under specific-pathogen-free conditions in the Experimental Biomedicine Unit at the University of Jena, Germany. Mice were bred according to registration number 02-053/16, and all experimental procedures were approved by the federal Thuringian “Landesamt für Verbraucherschutz” under registration number 02-030/15. All legal specifications were followed regarding European guidelines 2010/63/EU. Mice were randomly assigned to different treatment groups and sacrificed by cervical dislocation or CO2 inhalation. For intraperitoneal injections, M-100 was solved in 7.5 % N-methyl-2-pyrrolidone (Carl Roth, Karlsruhe, Germany, Cat#P052) and 40 % PEG-400 (Carl-Roth, Cat#0144) in sterile water.

### Cell lines and primary cultures

All cell lines were maintained in incubators at 37 °C and 5 % CO2. 293T and NIH-3T3 cells were grown in DMEM with 10 % FCS (Sigma-Aldrich, St. Louis, MO, United States, Cat#F7524). Ramos, Raji, and MV4-11 cells were grown in RPMI 1640 with 10 % FCS. BL-30 cells were grown in RPMI 1640 with 20 % FCS. OCI-Ly3, SUDHL-6, and CH12F3 cells were grown in RPMI 1640, supplemented with 10 % FCS, 50 μM β-mercaptoethanol, 10 mM HEPES. CH12F3 cells were activated by stimulation with 1 μg/ml CD40L (Thermo Fisher Scientific, Waltham, MA, United States, Cat#16-0402-81, RRID:AB_468944), 5 ng/ml IL-4 (Thermo Fisher Scientific, Cat#14-8041-62) and 1 ng/ml TGF-β1 (Cell Signaling Technology, Danvers, MA, United States, Cat#8915). All cell lines were tested regularly for *Mycoplasma* infection. Isolation of primary B-cells was performed as previously described (56). Primary B-cells were grown in RPMI 1640 with 10 % FCS, 50 μM β-mercaptoethanol, 10 mM HEPES and 0.5 % gentamicin. Activation of primary B-cells was induced by the addition of 10 μg/ml LPS (from *E. coli*, O111:B4, Sigma-Aldrich, Cat#L4391). Ramos cells were authenticated by Eurofins Genomics (Ebersberg, Germany) using PCR-single-locus-technology with 21 independent PCR-systems. M-100 was dissolved to a 10 mM stock solution with DMSO and diluted to a concentration of 100 μM with PBS. M-100 is registered under patent number WO2016020369 A1. Venetoclax (Selleck Chemicals, Houston, TX, United States) was solved in DMSO.

### Flow cytometry

All measurements were performed using a LSR Fortessa system (BD Biosciences, Franklin Lakes, NJ, United States), and data were acquired with BD FACSDIVA V8.0.1 (BD Biosciences). Immune cell phenotyping was performed by staining single-cell suspensions in PBS with respective antibodies. For apoptosis detection, an Annexin V-FITC Apoptosis Detection Kit (Thermo Fisher Scientific, RRID:AB_2575600) was used. Cell cycle analysis was performed with fixed cells by propidium iodide (PI) incorporation. Flow cytometry data were analyzed with FlowLogic 700.2A (Inivai Technologies, Mentone, Australia).

### Immunoprecipitation

Protein levels were determined using the Roti-Nanoquant solution (Carl Roth GmbH, Cat#K880). For IPs, lysates containing 250 μg (overexpressed) or 1000 μg (endogenous) protein were combined with a mixture of 50 % (v/v) protein A and 50 % (v/v) protein G beads (Sigma-Aldrich Inc., Cat#P9424 and Cat#P3296) and 0.5-1 μg antibody in lysis buffer overnight at 4 °C. Following control antibodies were used: Mouse IgG control (Santa Cruz Biotechnology, Dallas, TX, United States, Cat#sc-2025, RRID:AB_737182), Rabbit IgG control (Santa Cruz Biotechnology, Cat#sc-2027, RRID:AB_737197). Beads were washed three times in lysis buffer after incubation, resuspended in 2x Laemmli buffer to a final 1x concentration, and boiled for 5 min at 95 °C.

### Proximity ligation assay

PLA was performed using Duolink In Situ PLA Reagents Red (Sigma-Aldrich, Cat#DUO92008) according to the manufacturer’s protocol. Ramos cells were attached to coverslips with 0.1 % poly-L-lysine (Sigma-Aldrich, Cat#P8920) for 1 h at RT. Cells were permeabilized, fixed with methanol for 10 min at −20 °C, blocked with Duolink blocking solution, and incubated with primary antibodies. PLA Probe Anti-Mouse MINUS (Sigma-Aldrich) and PLA Probe Anti-Rabbit PLUS (Sigma-Aldrich) were used as secondary probes. Samples were mounted with DAPI. PLA signals were detected using a Nikon Ti Microscope (λex=594 nm; λem=624 nm). Images were taken with a Nikon DS-Qi2 camera. PLA foci were counted automatically with ImageJ (57) after threshold adjustment using function “analyze particles”.

### Proteomics

Immunoprecipitated proteins were eluted by the addition of NuPAGE LDS sample buffer (Thermo Fisher Scientific) supplemented with 1 mM DTT. The samples were heated at 70 °C for 10 min, alkylated with 5.5 mM chloroacetamide for 30 min in the dark, and loaded on NuPAGE 4-12 % gradient Bis-Tris gels (Thermo Fisher Scientific). Proteins were stained with Colloidal Blue and digested in-gel using trypsin. Peptides were extracted from the gel and desalted on reversed-phase C18 StageTips. Peptide fractions were analyzed on a quadrupole Orbitrap mass spectrometer (Q Exactive Plus, Thermo Fisher Scientific) equipped with a UHPLC system (EASY-nLC 1000, Thermo Fisher Scientific). Raw data files were analyzed using MaxQuant version 1.5.2.8 (58). Parent ion and MS2 spectra were searched against a UniProtKB database using the Andromeda search engine (59). Spectra were searched with a mass tolerance of 6 ppm in MS mode, 20 ppm in HCD MS2 mode, strict trypsin specificity, and allowing up to three miscleavages. Cysteine carbamidomethylation was searched as a fixed modification, whereas protein N-terminal acetylation and methionine oxidation were searched as variable modifications. Functional annotation clustering was performed with the DAVID tool (60).

## Acknowledgments

We thank Carmen Mertens for technical assistance as well as Dr. Karl-Gunther Glowalla, Petra Grübner and Andreas Köber for animal care. We are grateful to Katrin Schubert for assistance with cell sorting (FLI Jena) and Dr. David Corujo for bioinformatic support. We also want to thank Dr. Marcus Buschbeck and Dr. Jeannine Diesch for giving feedback on the manuscript. Prof. Dr. Ralf Küppers kindly provided lymphoma cell lines. Venetoclax was a kind gift from Prof. Dr. Sebastian Scholl and Dr. Maximilian Fleischmann.

## Authorship Contributions

Conception and design, C.K.; Development of methodology, R.W., A.-S.M., M.B., M.E.H., S.P., and F.H.; Acquisition of data, R.W., A.-S.M., E.-M.P., M.K., M.B., K.L., L.H., A.-M.S., M.E.H., S.P., and F.H.; Analysis and interpretation of data, R.W., A.-S.M., E.-M.P, M.E.H., and C. K.; Writing, review, and/or revision of the manuscript, R.W., A.-S.M., and C. K.; Administrative, technical, or material support, O.H.K., P.B., and S.M.; Study supervision, P.B., T.M, O.H.K., and C. K.

