## Supplemental Material for "Targeting the MYC interaction network in B-cell lymphoma via histone deacetylase 6 inhibition"

### Supplemental Figures

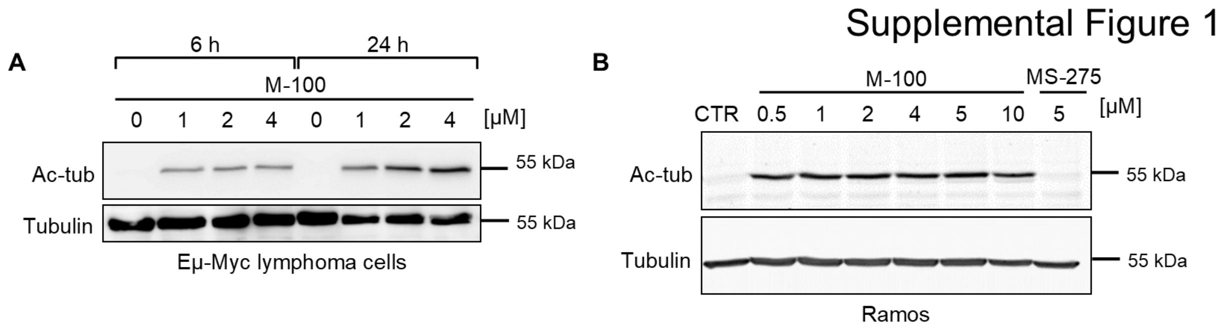


**Supplemental Figure 1: M-100 has no effect on unmodified tubulin levels.**

**(A)** Murine Eµ-Myc lymphoma and **(B)** human Ramos cells were treated with different concentrations of M-100 for 24 h. MS-275 treatment serves as negative control. Shown is one representative Western blot from n=2 independent experiments each.

Supplemental Figure 2

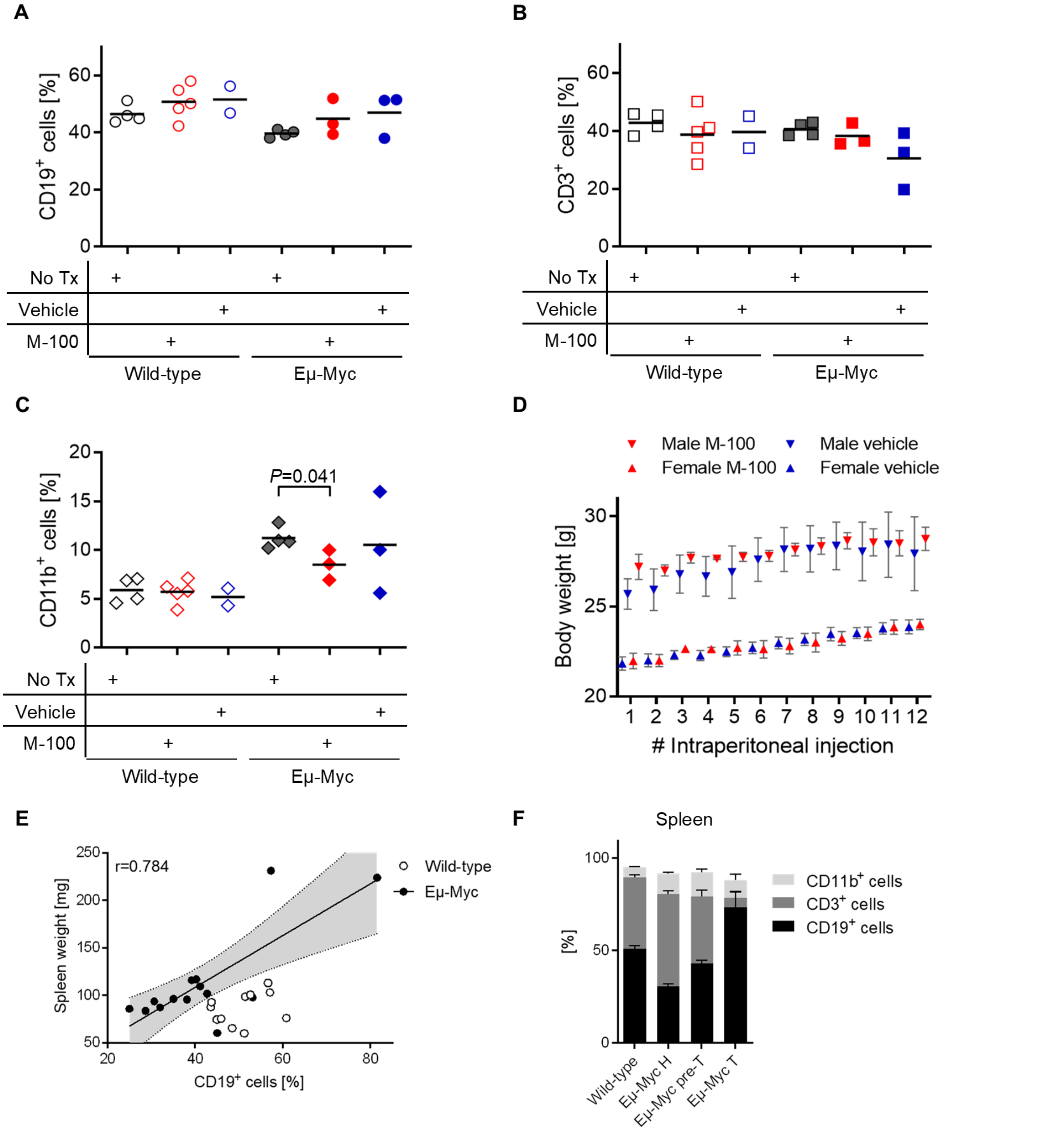


**Supplemental Figure 2: Short-term and long-term effects of M-100 *in vivo*.**

**(A-C)** Immune cell populations were analyzed in the spleens of 70-day-old wild-type and Eμ-Myc mice treated once via i.p. injection with 30 mg/kg M-100 or vehicle for 72 h by flow cytometry. (A) B-cells, (B) T-cells and (C) myeloid cells. Cell populations were compared to age-matched untreated mice. Each dot represents one individual mouse. Bars depict mean. One-Way ANOVA (Tukey's posthoc). **(D)** Body weights were measured of male and female Eμ-Myc mice treated via i.p. injection with M-100 (30 mg/kg) or vehicle for 12 times. Data represent mean ± SEM for each sex, treatment group, and number of i.p. injection. **(E)** Correlation between percentage of splenic $CD19^+$ cells and spleen weight of wild-type and Eμ-Myc mice aged 3 to 6 months. Each dot represents one individual mouse. Linear trend with 95 % confidence interval was inserted and Pearson's r was calculated for Eμ-Myc mice. **(F)** Eμ-Myc mice were grouped based on percentage of splenic $CD19^+$ cells into healthy (H, ≤35 %), pre-tumor (pre-T, ≤55 %), and tumor (T) stages. Shown are corresponding percentages of splenic $CD3^+$ and $CD11b^+$ cells. Bars depict mean + SEM from 11 (wild-type), 5 (H), 7 (pre-T), and 5 (T) individual mice per group.

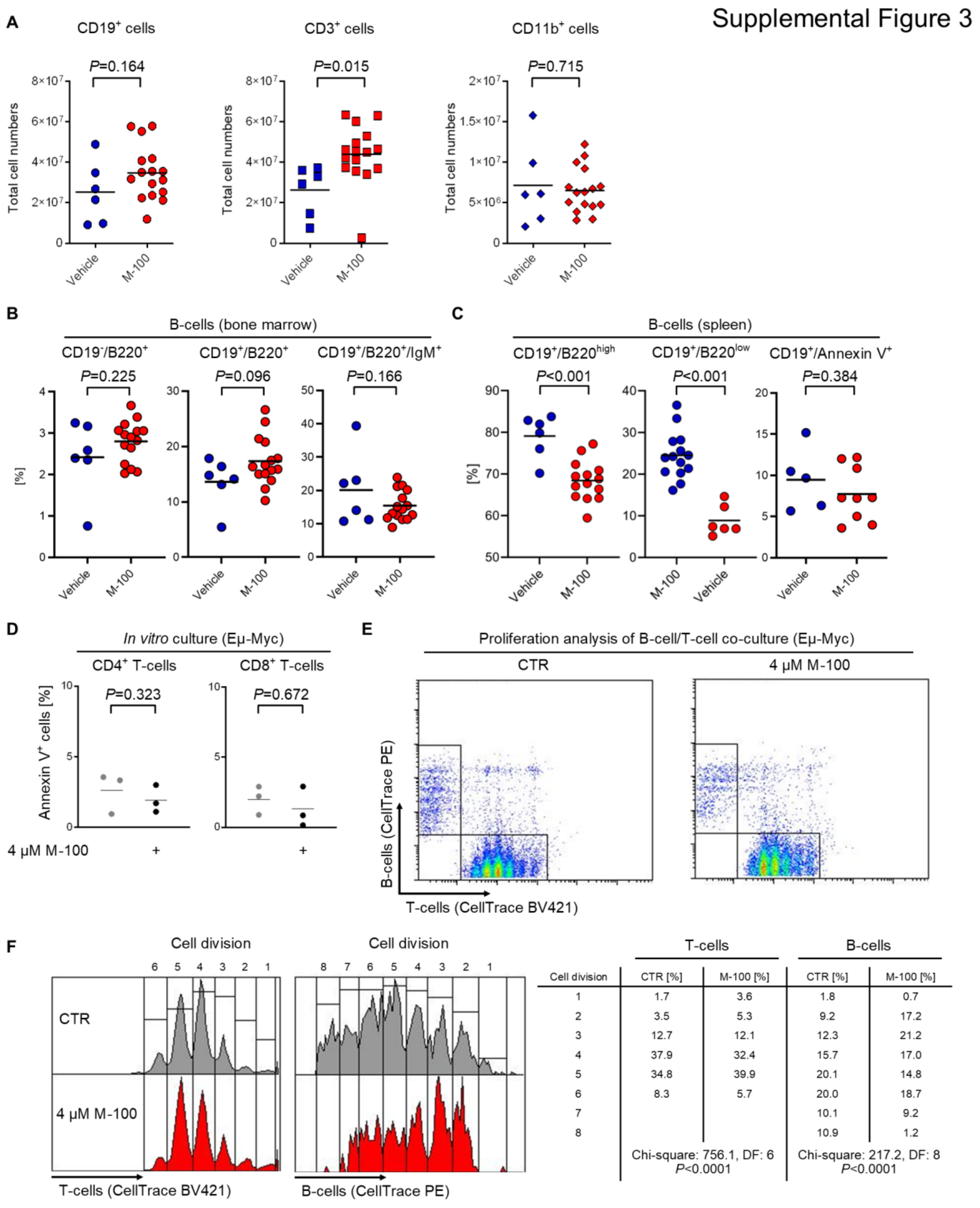


| | T-cells | | B-cells | |
|---|---|---|---|---|
| Cell division | CTR [%] | M-100 [%] | CTR [%] | M-100 [%] |
| 1 | 1.7 | 3.6 | 1.8 | 0.7 |
| 2 | 3.5 | 5.3 | 9.2 | 17.2 |
| 3 | 12.7 | 12.1 | 12.3 | 21.2 |
| 4 | 37.9 | 32.4 | 15.7 | 17.0 |
| 5 | 34.8 | 39.9 | 20.1 | 14.8 |
| 6 | 8.3 | 5.7 | 20.0 | 18.7 |
| 7 | | | 10.1 | 9.2 |
| 8 | | | 10.9 | 1.2 |
| | Chi-square: 756.1, DF: 6 *P*<0.0001 | | Chi-square: 217.2, DF: 8 *P*<0.0001 | |

**Supplemental Figure 3: Extended analysis of Eμ-Myc mice and immune cells treated with M-100.**

**(A)** Absolute cell counts for immune cell populations shown in **Figure 1F**. Unpaired Student's t-test, two-tailed. **(B)** Flow cytometry analysis of B-cells in the bone marrow from survivors. Each dot represents one mouse. Unpaired Student's t-test, two-tailed. **(C)** Flow cytometry analysis of $CD19^+$ B-cells in the spleen from survivors. $B220^{high}$ cells correspond to mature B-cells while $B220^{low}$ B-cells are considered pre-malignant. Apoptosis of splenic B-cells after long-term treatment with M-100 was analyzed by staining for Annexin V. Unpaired Student's t-test, two-tailed. **(D)** $CD4^+$ and $CD8^+$ T-cells were isolated from Eµ-Myc mice by positive selection, activated *in vitro* with 2 µg/ml immobilized CD3 and 2 µg/ml soluble CD28 for 24 h, and then treated with 4 µM M-100 for 48 h. Apoptosis was analyzed by Annexin V staining. Each dot represents cells from one mouse. Bars depict mean. Paired Student's t-test, two-tailed. **(E)** B-cells and $CD4^+$ T-cells were isolated from Eµ-Myc mice, stained with different proliferation dyes, and co-cultured together. T-cells were activated as described above and 4 µM M-100 were added 24 h after activation. Proliferation of B-cells and T-cells was analyzed by flow cytometry 48 h after M-100 treatment. Loss of fluorescence intensity corresponds to cell divisions. **(F)** Proliferation from (E) was analyzed by gating on each cell division for B-cells and T-cells. Distributions were compared using Chi-square test. DF - degrees of freedom. All bars depict mean.

Supplemental Figure 4

**A**

| **Grade** | **Activity** | **Tumors** | **Posture** | **Observations** |
|---|---|---|---|---|
| 1: none | Active, strong | None | Normal | None |
| 2: low | Less active, with interrup-tions | Small lymph node lym-phomas | Slightly hunched | Weight +/- 10 % |
| 3: mid | Slow, sleepy, moves with difficulty | Additional lymphomas in cervical region | Hunched, trembling | Weight +/- 15 %, bloat-ed stomach, furry |
| 4: high | Lethargic, motionless | Lymphoma diameter >1 cm | Severely hunched, trembling | Moribund, diffi-culties to breath |

**B**

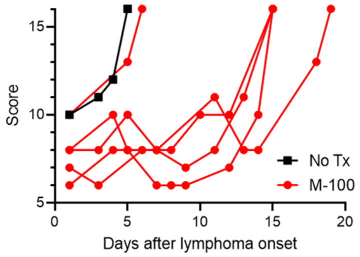


**Supplemental Figure 4: Score sheet for assessing disease severity.**

**(A)** Score table. The final score is calculated by adding points achieved in each category.

**(B)** Measured scores over time for Eμ-Myc animals bearing lymphomas are shown. The maximum score was given for animals reaching endpoint criteria or death.

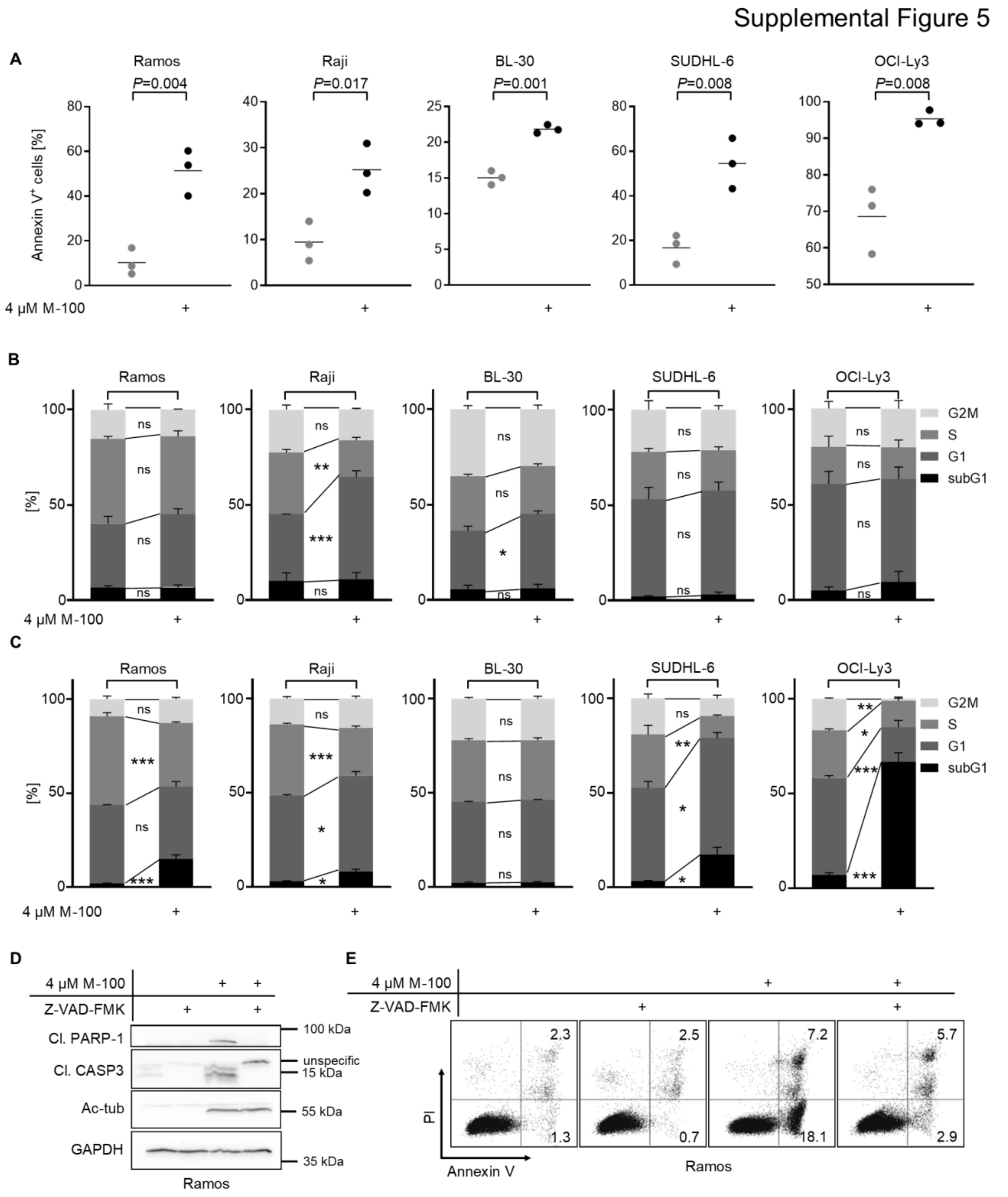


**Supplemental Figure 5: HDAC6 inhibition induces apoptosis in human B-cell lymphoma cell lines.**

**(A)** Amount of Annexin V$^{+}$ cells after treatment with 4 µM M-100 for 72 h. Unpaired

Student's t-test, two-tailed. **(B, C)** Cell cycle analysis of B-cell lymphoma cell lines by PI staining. Cells were left untreated or treated for (B) 24 h or (C) 72 h with 4 µM M-100. Two-Way ANOVA (Sidak's posthoc). **(D)** Western Blot analysis of Ramos cells treated for 24 h with 4 µM M-100 and 100 µM Z-VAD-FMK to block caspase activity. GAPDH was used as a loading control. Ac-tub - acetylated tubulin, cl. - cleaved. **(E)** Annexin V and PI staining of Ramos cells treated for 48 h as in (D). Data in (A) - (C) and (E) are representative of n=3, and data in (D) are representative of n=2 independent experiments. Data in (A) - (C) represent mean + SEM. *$P<0.05$, **$P<0.01$, ***$P<0.001$, ns - not significant.

Supplemental Figure 6

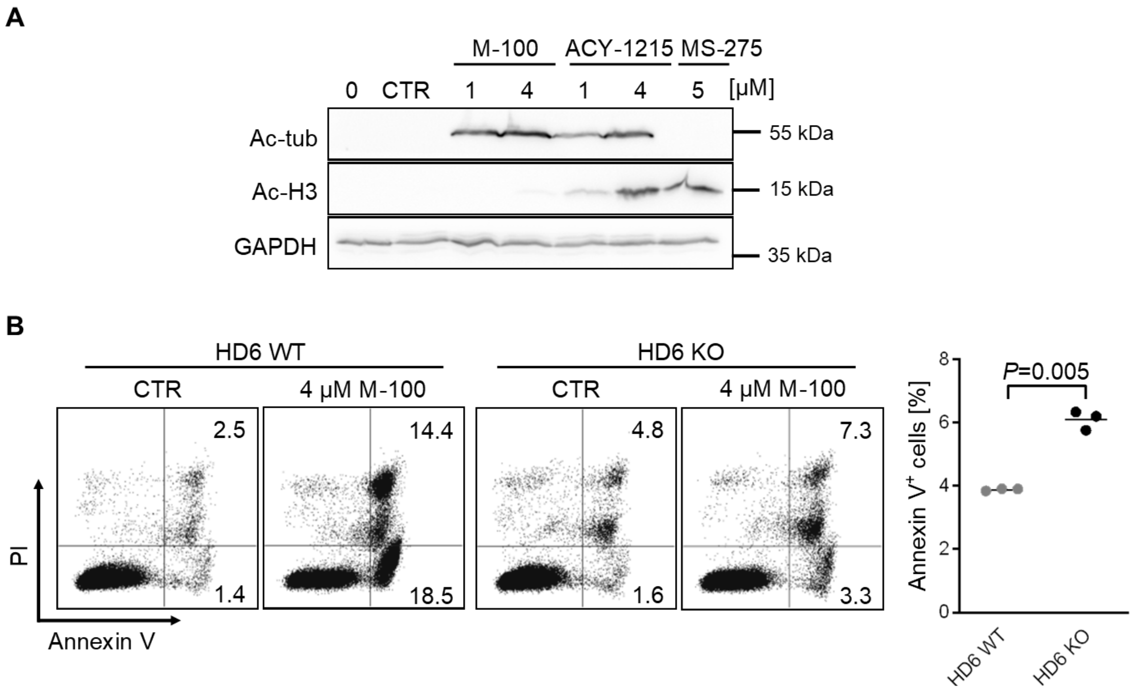


**Supplemental Figure 6: M-100 is a highly specific HDAC6 inhibitor.**

**(A)** Western blot analysis of off-target effects for M-100 and ACY-1215 by comparing acetylated tubulin (ac-tub) and acetylated histone 3 (ac-H3) signals. Treatment with pan-HDACi MS-275 serves as a control for ac-H3 signals. All treatments were performed for 24 h. GAPDH was used as a loading control. **(B)** Apoptosis was analyzed by Annexin V/PI staining in Ramos HDAC6 WT and Ramos HDAC6 KO cells treated for 72 h with 4 µM M-100 or left untreated. Percentages of Annexin $V^+$ cells were compared for untreated conditions. Bars depict mean. Data represent mean + SEM. Unpaired Welch's t-test, two-tailed. Data in (A) and (B) are representative of at least n=3 independent experiments.

Supplemental Figure 7

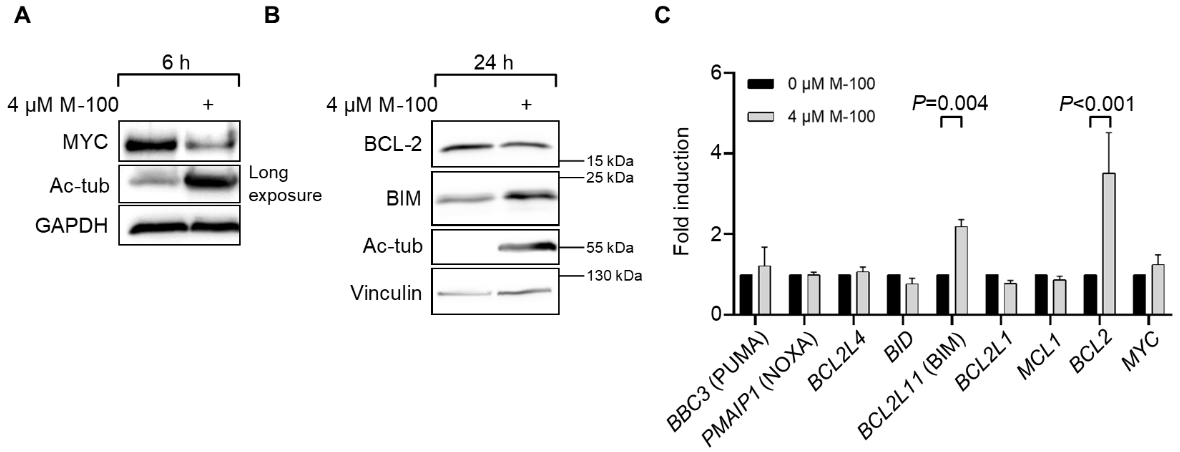


**Supplemental Figure 7: M-100 affects expression of genes related to apoptosis.**

**(A, B)** Western blot analysis of Ramos cells treated for (A) 6 h or (B) 24 h with 4 µM M-100. GAPDH or Vinculin serve as loading control. **(C)** Gene expression changes of Ramos cells treated for 24 h with M-100 compared to untreated cells using quantitative real-time PCR analysis. Data represent mean + SEM. Multiple t-test.

Data in (A) and (B) are representative of n=2 independent experiments and data in (C) are representative of at least n=3 independent experiments.

Supplemental Figure 8

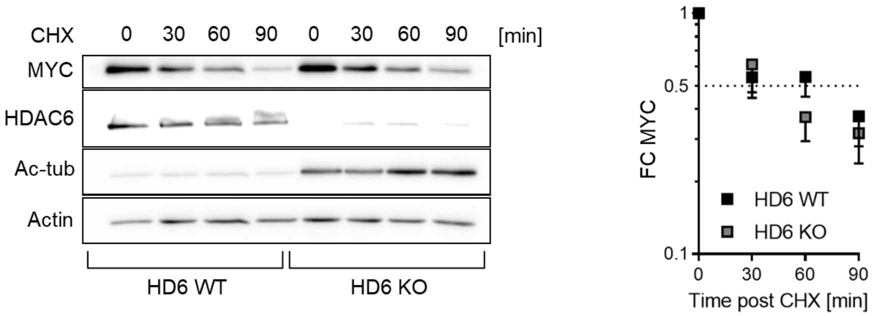


**Supplemental Figure 8: Knock-out of HDAC6 has minor effects on MYC turnover.**

Western Blot analysis was performed of Ramos HDAC6 WT (HD6 WT) and KO cells treated for the indicated time points with cycloheximide (CHX, 50 µg/ml). Actin serves as loading control. Shown is one representative Western blot from n=4 independent experiments. MYC levels were quantified and normalized. Data represent mean - SEM. Ac - acetylated, FC - fold change.

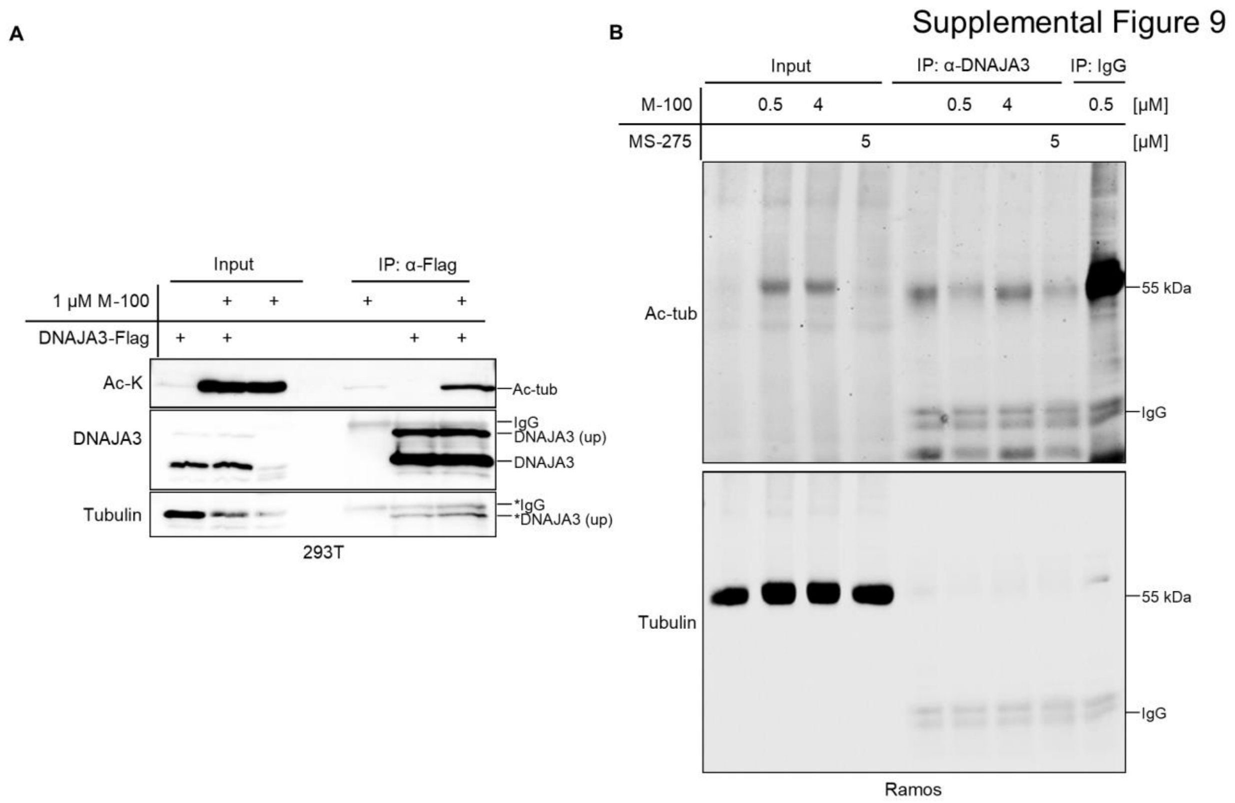


**Supplemental Figure 9: Unmodified tubulin does not associate with DNAJA3.**

**(A)** Samples shown in **Figure 6A** were tested for unmodified tubulin. **(B)** Experiment was performed as in **Figure 6B**. Ramos cells were treated for 24 h with either 0.5 µM, 4 µM M-100, 5 µM MS-275, or left untreated. Lysates were used for IP with α-DNAJA3 antibodies and tested for interaction with unmodified tubulin. IPs with unspecific IgG were used as control.

Supplemental Figure 10

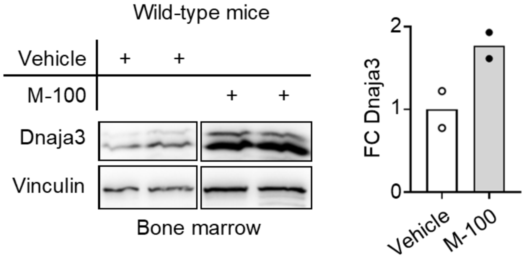


**Supplemental Figure 10: Dnaja3 is increased in wild-type mice after M-100 treatment.**

Western Blot analysis was performed of bone marrow lysates from wild-type mice after one i.p. injection with M-100 (30 mg/kg) or vehicle. Small and large isoforms of Dnaja3 can be noticed. Vinculin was used as a loading control. Each lane represents one individual mouse. Quantification of Dnaja3 protein levels is shown based on Vinculin. FC - fold change.

### Supplemental Tables

**Supplemental Table 1: Identified proteins from mass spectrometry.** Please refer to the Excel file.

### Supplemental Methods

#### Transfection and cell lysis

The medium was changed to medium without FCS before the transfection of adherent cells. Following recombinant DNAs were used: DNAJA3-Flag (created by us), pcDNA3.1 (Thermo Fisher Scientific Inc., Cat#V79020), pcDNA3.1 MYC (1), pCMV hTid Long Addgene Plasmid #13707 (2), pCMV hTid Short Addgene Plasmid #13709 (2), pEGFP C2 (Clontech), pEGFP C2 MYC (created by us). For 500 000 cells, DNA was mixed with control plasmids up to 4 µg DNA in reaction tubes containing 120 µl PBS. 10.8 µl polyethyleneimine (10 mM, Sigma-Aldrich Inc., Cat#408727) was diluted in 120 µl PBS and then mixed with DNA. Transfection complexes were added to cells after incubation for 20 min at RT, and the medium was changed to complete medium 6 h post-transfection. Cells were lysed 48 h post-transfection in cell lysis buffer: 50 mM Tris-HCl pH 8.0, 150 mM NaCl, 1 % (v/v) Nonidet P-40, 0.2 % (v/v) protease inhibitors: 1 mg/ml leupeptin (Carl Roth GmbH, Cat#CN33.1), 2 mg/ml antipain (Carl Roth GmbH, Cat#2933.2), 100 mg/ml benzamidine (Sigma-Aldrich Inc., Cat#B6506), 10 kU/ml aprotinin (Carl Roth GmbH, Cat#A162) or stringent lysis buffer (lysis buffer supplemented with 1 mM EDTA pH 8.0, 1 mM EGTA pH 8.0, 1 % (w/v) sodium deoxycholate, 0.1 % (w/v) SDS) and incubated for 20 min on ice. Lysates were sonicated for 5 min using a water-bath sonicator (Sonorex Super RK106, Bandelin) and centrifuged at 13 000 g for 10 min.

#### SDS-PAGE and Western blot

Proteins were mixed with 6x Laemmli buffer (35 % (v/v) β-mercaptoethanol, 350 mM Tris-HCl pH 6.8, 30 % (v/v) glycerin, 10 % (w/v) SDS, 0.25 % (w/v) bromophenol blue) to a final 1x concentration and boiled for 5 min at 95 °C. Proteins were separated on freshly prepared 7-12 % (v/v) polyacrylamide separating gels with 5 % (v/v) polyacrylamide stacking gels at 80 V for approximately 2 h. PageRuler Plus Prestained Protein Ladder (Thermo Fisher Scientific Inc., Cat#26619) was included as a reference in the run. Proteins were transferred onto ethanol-activated PVDF membranes (Carl Roth GmbH Cat#T830) using 1x Towbin buffer (25 mM Tris, 250 mM glycine), placed between sheets of chromatography paper (GE Healthcare, Chicago, IL, United States, Cat#3030-347). Blotting was performed at 150 mA per gel for 2 h or at 20 mA per gel overnight. Non-specific bindings were blocked using 5 % (w/v) non-fat dry milk (Saliter) in PBS (10x PBS: 1.37 M NaCl, 27 mM KCl, 80 mM, $Na_2HPO_4$ x 2 $H_2O$, 14 mM $KH_2PO_4$) with 0.05 % (v/v) Tween 20 for 1 h at RT. Membranes were incubated with primary antibodies in 2 % (w/v) non-fat dry milk in PBS-Tween 20 according to the manufacturer's recommendation at 4° C overnight. Membranes were washed on the next day three times in PBS-Tween 20 for 5 min and incubated with corresponding HRP-conjugated secondary antibodies (Thermo Fisher Scientific Inc., Cat#31430, RRID:AB_228307 and Cat#31460, RRID:AB_228341) for 1 h at RT. Protein visualization was performed using ECL Western Blotting Substrates (ECL Westar Etac Ultra 2.0, Cyanagen Srl, Bologna, Italy, Cat#XLS0750100, or ECL Western Blotting Substrate, Thermo Fisher Scientific Inc., Cat#32106) with the Fusion Solo 4S system (Vilber Lourmat, Eberhardzell, Germany) and FusionCapt Advanced 17.03 software (Vilber Lourmat). Stepwise detection was performed of proteins using primary antibodies from different producer species on the same membrane. Quantification of Western Blots was performed with BIO-1D advanced software (Vilber Lourmat).

**Cell fractionation**

Cells were washed twice in PBS and centrifuged for 5 min at 700 g, RT. Then, pellets were resuspended in ice-cold cytoplasma extraction buffer (10 mM HEPES-KOH, pH 7.6, 15 mM KCl, 2 mM $MgCl_2$, 0.1 mM EDTA, freshly prepared 1 mM DTT, protease inhibitors) and incubated for 10 min on ice. Suspensions were centrifuged for 5 min at 700 g, RT, and supernatant (cytoplasmic extract) was collected. Residual pellets were washed twice in PBS and centrifuged for 5 min at 700 g, RT. Pellets were resuspended in ice-cold nuclear extraction buffer (20 mM HEPES-KOH, pH 7.9, 420 mM NaCl, 2 mM $MgCl_2$, 0.2 mM EDTA, 25 % (v/v) glycerin, 1 mM DTT; protease inhibitors) and incubated for 15 min at 4 °C. The supernatant (nuclear extract) was transferred into new reaction tubes after centrifugation for 10 min at 10000g and sonicated for 5 min using a water-bath sonicator (SONOREX SUPER RK106, Bandelin).

**Proliferation analysis**

Primary splenic T-cells were purified by positive selection with anti-mouse CD4 biotin antibody (Miltenyi Biotech, Bergisch-Gladbach, Germany, Cat#130-101-962, RRID:AB_2659917) or CD8 biotin antibody (Miltenyi Biotech, Cat#130-118-074, RRID:AB_2733537) and anti-biotin MicroBeads (Miltenyi Biotech, Cat#130-090-485). B-cells were stained with CellTrace™ Yellow Cell Proliferation Kit and and T-cells with CellTrace™ Violet Cell Proliferation Kit (Invitrogen, Cat#C34567 and Cat#C34557), respectively, according to manufacturer's instruction. Then, B- and T-cells were seeded at a 1:1 ratio in primary B-cell medium. T-cells were activated with 2 µg/ml immobilized CD3 (BioLegend, San Diego, CA, Cat#100238) and 2 µg/ml soluble CD28 (BioLegend, Cat#102116). M-100 was added to the co-culture 24 h after T-cell activation for 48 h.

### Antibodies

Following primary antibodies were used: Mouse Anti-acetyl Tubulin, alpha, Monoclonal Antibody K40, Unconjugated. Clone 6-11B-1 (Santa Cruz Biotechnology, Cat#sc-23950, RRID:AB_628409); Mouse Anti-acetyl-Tubulin Monoclonal Antibody, Unconjugated, Clone 6-11B-1 (Sigma-Aldrich Inc., Cat#T5168, RRID:AB_477579); Mouse Anti-FLAG M2 Monoclonal Antibody, Unconjugated (Sigma-Aldrich Inc., Cat#F3165, RRID:AB_259529); Mouse Anti-HDAC1 Monoclonal Antibody, Unconjugated, Clone 10E2 (Santa Cruz Biotechnology, Cat#sc-81598, RRID:AB_2118083); Mouse Anti-Human Vinculin Monoclonal Antibody, Unconjugated, Clone V284 (Bio-Rad/AbD Serotec, Cat#MCA465S, RRID:AB_2214389); Mouse Anti-PARP1, Cleaved Form Monoclonal Antibody, Unconjugated, Clone F21-852 (BD Biosciences Inc., Cat#552596, RRID:AB_394437); Mouse Anti-TID1 Monoclonal Antibody, Unconjugated, Clone RS13 (Abcam Ltd., Cambridge, United Kingdom, Cat#ab81700, RRID:AB_1640415); Rabbit Anti-acetyl Histone H3 Polyclonal Antibody, Unconjugated (Millipore, Burlington, MA, United States, Cat#06-599, RRID:AB_2115283); Rabbit Anti-acetyl Lysine Polyclonal Antibody, Unconjugated (Millipore, Cat#ST1027-50UL, RRID:AB_10682447); Rabbit Anti-Actin Polyclonal Antibody, Unconjugated (Sigma-Aldrich Inc., Cat#A2066, RRID:AB_476693); Rabbit Anti-Bcl-2 Polyclonal antibody, Unconjugated, Clone N-19 (Santa Cruz Biotechnology, Cat#sc-492, RRID:AB_2064290); Rabbit Anti-Cleaved Caspase-3 (Asp175) Monoclonal Antibody, Unconjugated, Clone 5A1E (Cell Signaling Technology, Cat#9664, RRID:AB_2070042); Rabbit Anti-MYC Polyclonal Antibody, Unconjugated, Clone N262 (Santa Cruz Biotechnology, Cat#sc-764, RRID:AB_631276); Rabbit Anti-GAPDH Polyclonal Antibody, Unconjugated (Sigma-Aldrich Inc., Cat#G9545, RRID:AB_796208); Rabbit Anti-HDAC6 Monoclonal Antibody, Unconjugated, Clone

D2E5 (Cell Signaling Technology, Cat#7558, RRID:AB_10891804); Anti-Mouse CD11b Monoclonal Antibody, FITC Conjugated, Clone M1/70 (Thermo Fisher Scientific Inc., Cat#11-0112-71, RRID:AB_464933); Hamster Anti-CD3e Monoclonal Antibody, PE Conjugated, Clone 145-2C11 (Thermo Fisher Scientific Inc., Cat#12-0031-81, RRID:AB_465495); Rat Anti-B220 Monoclonal Antibody, APC Conjugated, Clone RA3-6B2 (Thermo Fisher Scientific Inc., Cat#17-0452-83, RRID:AB_469396); Rat Anti-CD19 Monoclonal Antibody, APC Conjugated, Clone 1D3 (BD Biosciences, Cat#550992, RRID:AB_398483); Rat Anti-CD4 Monoclonal Antibody, APC Conjugated, Clone RM4-5 (Thermo Fisher Scientific Inc., Cat#17-0042-82, RRID:AB_469323); Rat Anti-CD8a Monoclonal Antibody, APC Conjugated, Clone 53-6.7 (Thermo Fisher Scientific Inc., Cat#17-0081-82, RRID:AB_469335).

**RNA isolation**

High-quality RNA was isolated using the Direct-zol RNA Miniprep Kit (Zymo Research, Irvine, CA, United States, Cat#R2052). In short, cells were lysed in RNAPure peqGOLD (VWR International Ltd., Radnor, PA, United States, Cat#30-1010) and mixed with an equal volume of 95 % (v/v) ethanol. RNA was bound to columns, washed, and subject to on-column-digestion with DNAse I (30 U) for 15 min at RT according to the manufacturer's protocol. RNA was eluted in RNAse-free water. The purity of RNA was measured by absorption at λ=230 nm, λ=260 nm, and λ=280 nm using a photometer (VWR International Ltd., ND-1000).

**Quantitative real-time PCR**

For quantitative real-time PCR experiments, cDNA was generated from up to 1 µg RNA using First Strand cDNA Synthesis Kit (Thermo Fisher Scientific Inc., Cat#K1612) according to the manufacturer's protocol, and an equally mixed combination of oligo-(dT)18 and random hexamer primers. Runs were performed on a StepOnePlus Real-Time PCR system (Thermo Fisher Scientific Inc.) using StepOne Software v2.3 (Life Technologies). PowerUp SYBR Green Master Mix (Thermo Fisher Scientific Inc., Cat#A25778) was combined with specific primers (200 pmol) and 5 ng cDNAs for a single reaction in 96-well plates (MicroAmp Fast Optical 96-well reaction plate, Applied Biosystems, Waltham, MA, United States, Cat#4346906) sealed with MicroAmp Clear Adhesive Film (Applied Biosystems, Cat#4306311). Technical triplets and negative controls were prepared for each reaction. Polymerase started amplification after an initial denaturation step at 95 °C, and all annealing steps were performed at 60 °C. Melting curves were generated for each primer pair. Data analysis was performed using the comparative ΔΔCT method. Fold inductions were calculated using the comparative ΔΔCT method based on *Actin* or *GAPDH* expression.

Following primer pairs were used for murine transcripts 5’ >3’: *Actin*: CTAAGGCCAACCGTGAAAAG and ACCAGAGGCATACAGGGACA, *Bbc3*: ACGACCTCAACGCGCAGTACG and GAGGAGTCCCATGAAGAGATTG, *Bcl2*: CTGAACCGGCATCTG and GGGGCCATATAGTTCCACAAA, *Myc*: TTTGTCTATTTGGGGACAGTGTT and CATCGTCGTGGCTGTCTG, *Pmaip1*: CAGATGCCTGGGAAGTCG and TGAGCACACTCGTCCTTCAA.

Following primer pairs were used for human transcripts 5’ ->3’: *BBC3*: CACCCTGGAGGGTCCTGTA and GCACCTAATTGGGCTCCATCT, *BCL2:* GGATCCAGGATAACGGAGGC and GAAATCAAACAGAGGCCGCA, *BCL2L1:* GTGAGTCGGATCGCAGCTT and GCTGCTGCATTGTTCCCATAG, *BCL2L11:*

CATCGCGGTATTCGGTTC and GCTTTGCCATTTGGTCTTTTT, *BCL2L4:* GTCTTTTTCCGAGTGGCAGC and TAGAAAAGGGCGACAACCCG, *BID:* TGCAGCTCAGGAACACCA and TCTCCATGTCTCTAGGGTAGGC, *GAPDH:* TGCACCACCAACTGCTTAGC and GGCATGGACTGTGGTCATGAG, *MCL1:* GGACAAAACGGGACTGGCTA and CAGCAGCACATTCCTGATGC, *MYC:* CACCAGCAGCGACTCTGA and CTGTGAGGAGGTTTGCTGTG, *PMAIP1:* CAGCTGTCCGAGGTGCTC and CCGCCCACTCAGCTACAG.

**CRISPR/Cas9-mediated deletion of *HDAC6***

The CRISPR/Cas9-system was utilized to delete *HDAC6* in Ramos cells with the following crRNA: *GCCGGUUGAGGUCAUAGUUGGUUUUAGAGCUAUGCU*. Oligonucleotides Alt-R CRISPR-Cas9 tracrRNA, ATTO 550 (Integrated DNA Technologies, Coralville, IA, United States, Cat#1075927) and Alt-R CRISPR-Cas9 crRNA (Integrated DNA Technologies) were mixed to a final duplex concentration of 100 µM, heated for 5 min at 95 °C and cooled down to RT. The formation of ribonucleoprotein complexes was achieved by mixing 120 pmol RNA duplex and 104 pmol Alt-R S. p. Cas9 Nuclease 3NLS (Integrated DNA Technologies, Cat# 1074181) with PBS to a final volume of 5 µl. The mixture was incubated for 20 min at RT and electroporated into Ramos cells using Cell Line Nucleofector Kit V (Lonza, Basel, Switzerland, Cat#VACA-1003) according to the manufacturer's protocol. Ribonucleoprotein complexes and Alt-R Cas9 Electroporation Enhancer (Integrated DNA Technologies, Cat#1075915) were mixed with cells, transferred to a cuvette and electroporation was performed in an Amaxa Nucleofector device (Lonza, program 0-06). Electroporated cells were resuspended in pre-warmed medium and cultivated. After 24 h, dead cells were eliminated using the Dead Cell

Removal Kit (Miltenyi Biotec, Cat#130-090-101) according to the manufacturer's protocol. Electroporated cells were sorted after 24 h using ATTO 550 label (BD FACSAria III, BD Biosciences), and expanded as single-cell clones in 96-well plates. Proteins were separated with SDS-PAGE and analyzed using Western Blot to check for genomic deletion. The genomic region of interest was amplified by conventional PCR and sequenced (Eurofins Genomics).
